# Catch*ReSeq resolves rare splice isoforms and their host cell states at single-cell resolution

**DOI:** 10.64898/2026.11.12.763766

**Authors:** Emanuelle I. Grody, Jinhee Kim, Kayla Yerlioglu, Amir Darbar, Gillian Primavera, Romaila Abdelraouf, Nicholas Baker, Jessica Derham, Diptatanu Das, Adunoluwa Obisesan, Leila B. Giron, Richard D’Aquila, Mohamed Abdel-Mohsen, Judd F. Hultquist, Ramon Lorenzo Redondo, Auinash Kalsotra, Elena Martinelli, Yogesh Goyal

## Abstract

Alternative splicing diversifies the transcriptome and shapes cell fate across development, tissue function, and disease. Single-cell RNA sequencing can link isoforms to cell states, but these methods often lack the throughput and coverage to accurately call splice variants. As a result, rare isoforms are often missed, especially in underrepresented cell types. Here, we present Catch*ReSeq, a targeted enrichment method that yields isoform-resolved readouts linked to host transcriptomes from standard single-cell libraries for the capture of rare isoforms. We found more oligo(dT)-primer cDNA originating from internal A-rich sequences than from terminal polyA tails (80-fold enrichment for simian immunodeficiency virus (SIV) and over 6-fold for host transcripts), fracturing cDNA and obscuring isoforms. Catch*ReSeq exploits this fracture to infer each molecule’s isoform of origin. In SIV-infected macaque CD4⁺ T cells, Catch*ReSeq detected up to 13 times as many infected cells as standard 10X and resolved spliced viral isoforms that 10X missed entirely. Splice+ cells carried a translational host program independent of how much virus they transcribed, a distinction invisible to standard single-cell sequencing methods. From macaque and PWH on ART, Catch*ReSeq recovered 13-fold more virus-transcribing cells than 10X, including splice+ cells. We also demonstrated the generalizability of Catch*ReSeq across biological systems by applying it in combination with lineage barcoding to probe alternative splicing in clonally expanded CD4⁺ T cells, and separately in mouse liver regeneration. As Catch*ReSeq works with archived cDNA and can be redesigned for new targets after the fact, it allows rare isoform profiling on existing biobanked and clinical samples.

## INTRODUCTION

Rare transcripts, whether be they rarely expressed genes or low-abundance isoforms of common genes, can carry important biological significance, yet their detection remains a central challenge in transcriptome profiling. This is further conflated in cases where rare transcripts are present in rare or underrepresented cells, compounding the detection problem within and between cells^1^. One important source of rare transcripts is alternative splicing, which generates transcript variants from a pre-mRNA through variable inclusion of exons^2^. By determining which parts of genes are expressed into proteins, alternative splicing contributes to differential outcomes across development, tissue function, and disease^2–4^ and can vary between individual cells and within a tissue or lineage^5–8^. Examples of rare and consequential alternative splicing are found in neurodegeneration^9^, liver regeneration^10^, and several other states of human health and disease^2,4^.

Alternative splicing is a common way by which viruses maximize the coding potential of their size-limited genomes. For example, the ∼9 kb primary transcript of human immunodeficiency virus (HIV) and its non-human primate counterpart simian immunodeficiency virus (SIV) undergoes alternative splicing to produce over 50 RNA species encoding distinct viral proteins essential for viral replication^11,12^. The balance of viral RNA isoforms present in an infected cell is dynamic throughout viral replication and is associated with whether the cell supports productive infection or persistence in a latent state^13–16^. Given the rarity of infected cells in specimens from people with HIV (PWH), especially during suppressive antiretroviral therapy (ART), detection of rare viral spliced isoforms at single-cell resolution remains challenging. While methods have been devised to preferentially sort for infected cells^17^ or to boost rare transcript number by stimulating viral transcription^18^, it is unclear how these manipulations alter the underlying biological meaning.

Studying how rare events such as alternative splicing shape cell fate requires both sensitive detection of the transcript variant and broad quantification of the host cell transcriptome. While alternative splicing can be detected through single-cell RNA sequencing (scRNASeq), high throughput library preparation methods (e.g. 10X Chromium 3’ or 5’) capture only one end of the transcript and coverage in gene bodies may be sparse, limiting reliable detection of alternative splicing events. Methods such as Smart-seq3^19^ provide coverage across the transcript body but at much lower cell throughput. Even with full-transcript coverage, detection of rare transcripts remains unreliable due to transcript dropout and amplification noise introduced during mRNA capture, PCR amplification, and library preparation^20^. Among methods for single-cell mRNA isoform detection^8,21–24^, direct capture strategies—achieved via target-specific primer spike-in during reverse transcription or on-bead capture—can boost sensitivity for user-selected transcripts. For instance, a recent study^22^ added custom primers during reverse transcription to increase detection of lowly expressed HIV transcripts, including non-polyadenylated species, from infected cells. However, direct capture is limited to a user-selected panel without the ability to expand the target set post-hoc. Some methods are not designed for rare transcript targeting but can be used for this purpose. For example, Smart-seq3 can be paired with marker-based rare cell enrichment (e.g., FACS), but this approach inherently biases toward a preselected cell state(s) permissive of the rare phenotype and requires a surface marker that may not exist for the process of interest. Other methods for probe-based enrichment for DNA or RNA of rare cells (e.g., FISH-flow) for transcriptome profiling do not have explicit strategies to identify rare isoforms within these cells^26,27^.

To address the challenge of rare isoform capture, we developed an integrated experimental and computational method for targeted enrichment of rare transcript variants and joint host cell transcriptomic profiling which we call Catch*ReSeq (“catch and re-sequence”). Catch*ReSeq libraries can be created at scale, in parallel, and sequenced with a fraction of the reads recommended for a 10X library. We leveraged Catch*ReSeq to map viral isoforms during acute SIV infection *in vitro* and *in vivo* in non-human primates. We found that oligo(dT) captures internal adenine-rich sequences up to 80 times more often than the terminal polyA tail, fracturing viral transcripts. We turned this bias into an advantage: because fractured molecules terminate within the transcript body, the alignment position of an enriched product reports whether the targeted splice site was used. Catch*ReSeq recovered 56–89% more virus-transcribing cells than 10X and 275-fold more reads spanning viral splice junctions (965 versus 3.5 per 10,000 viral reads). Catch*ReSeq identified a translational host program enriched in splice+ cells independent of viral transcript load, a distinction inaccessible to 10X which detected predominantly high viral transcript load cells. Applied to rare infected cells from macaque and PWH on ART, Catch*ReSeq recovered 13-fold more virus-transcribing cells than 10X and identified an oxidative phosphorylation-depleted cell state permissive to viral transcription and splicing despite virologic control. We demonstrated the generalizability of Catch*ReSeq by applying it to probe alternative splicing in two other biological systems: lineage barcoded and clonally expanded CD4+ T cells, and liver regeneration. Ultimately, Catch*ReSeq’s compatibility with archived 10X 3’ cDNA libraries will enable targeted and high-throughput profiling of rare transcript isoforms across a wealth of existing samples.

## RESULTS

### Catch*ReSeq is specific and sensitive for targets

Catch*ReSeq design augments scRNAseq datasets with targeted splicing isoform resolution (**Fig 1A**, **Supp Fig 1A**). After cDNA amplification, only a quarter of the cDNA volume is used to create the 10X Gene Expression library (hereafter “10X”). Catch*ReSeq is performed with a subset of the remaining cDNA. First, we PCR amplify the cDNA to enrich for target regions by designing a forward primer upstream of the splice site or within the variably included exon and a reverse primer that binds to a common 10X handle. Enriched products then undergo 10X short-read library preparation. The reverse primer binds outside of the UMI and cell barcode, such that these labels are shared between both the Catch*ReSeq and 10X libraries for downstream analysis and minimizing PCR amplification artifacts (**Methods**). As part of our computation pipeline that joins Catch*ReSeq and full transcriptome datasets, we developed conservative quality control restrictions including optimized reads per UMI and UMIs per cell thresholds to remove false positives from PCR enrichment errors and spurious products (**Supp Fig 1A,B**; **Methods**).

**Figure 1:**
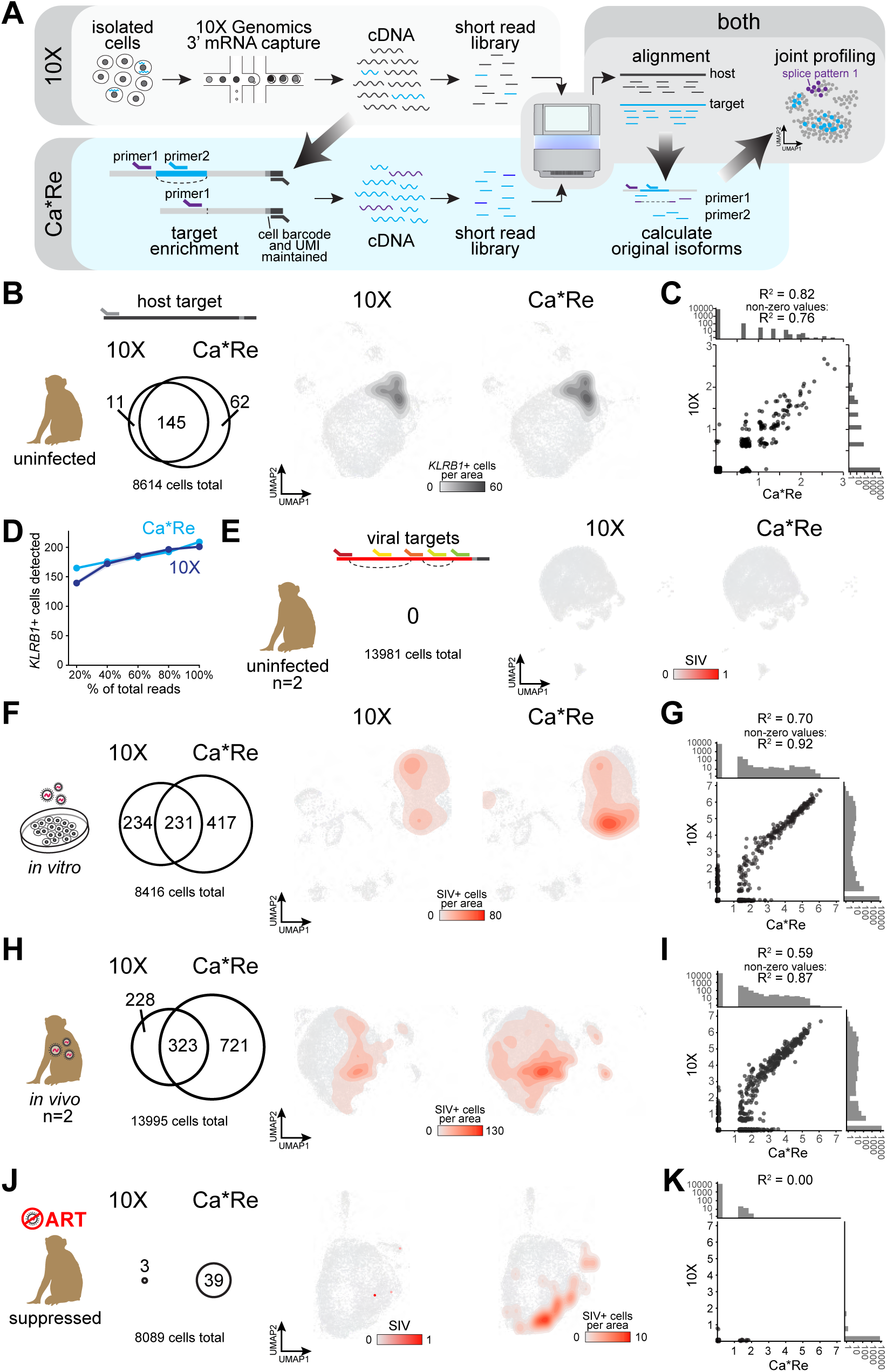
Sensitive capture of specific targets in host and viral mRNA. A. Schematic overview of Catch*ReSeq (“Ca*Re”) method. Catch*ReSeq applies custom targeted primers on cDNA from 10X Gene Expression (“10X”) protocol to generate an enriched short-read library. Sequencing and read alignment, Catch*ReSeq calculates the original isoforms based on primer used and read alignment, which is joined to the full transcriptome for joint profiling. B. *KLRB1*+ cell density of single cells captured by 10X (center) and Catch*ReSeq (right) in UMAP space (n = 8,614 cells). A Venn diagram (left) summarizes *KLRB1*+ cells per method: 11 10X-only, 145 shared, 62 Catch*ReSeq-only (218 positive total). Contour fill (dark gray) is the number of *KLRB1*+ cells per 1% of the UMAP bounding-box area. C. Correlation of *KLRB1* expression across single cells between 10X and Catch*ReSeq (n = 8,614 cells; Pearson R^2^ = 0.82 across all cells, R^2^ = 0.76 across 145 cells positive in both methods). Marginal histograms of cell counts are shown on a pseudo-log_10_ scale. D. Saturation analysis of *KLRB1*+ cells detected under binomial read subsampling (20–100% of total reads, 5 replicates per depth) for 10X (navy) and Catch*ReSeq (blue). Lines show the mean and shaded bands ±1 SD across replicates. A cell is counted positive at ≥1 read (10X) or ≥5 reads (Catch*ReSeq). E. SIV+ cell density of single cells from uninfected macaque cells captured by 10X (center) and Catch*ReSeq (right) in UMAP space (n = 13,981 cells; 0 SIV+ cells each). Counts are ln(x+1)-transformed and rescaled in arbitrary units (red = 1). F. SIV+ cell density of single cells in *in vitro*-infected macaque cells captured by 10X (center) and Catch*ReSeq (right) in UMAP space (n = 8,416 cells). Venn diagram (left): 234 10X-only, 231 shared, 417 Ca*Re-only. Contour fill (red) is the number of SIV+ cells per 1% of the UMAP bounding-box area. G. Correlation of SIV expression in single cells from (F) between 10X and Catch*ReSeq (n = 8,419 cells; Pearson R^2^ = 0.70 all cells, R^2^ = 0.92 across 231 both-positive cells). Marginal histograms shown on a pseudo-log_10_ scale. H. SIV+ cell density of single cells as captured by 10X (center) and Catch*ReSeq (right) from *in vivo* infected macaque cells during viremia from two individuals (integrated) in UMAP space. Venn diagram (left): 228 10X-only, 323 shared, 721 Ca*Re-only. Contour fill (red) is the number of SIV+ cells per 1% of the UMAP bounding-box area. I. Correlation of SIV expression in single cells from (H) between 10X and Catch*ReSeq (n = 13,995 cells; Pearson R^2^ = 0.59 all cells, R^2^ = 0.87 across 323 both-positive cells). Marginal histograms on a pseudo-log_10_ scale. J. SIV+ cell density of *in vivo*-infected macaque cells during ART suppression (D195) captured by 10X (center) and Catch*ReSeq (right) in UMAP space (n = 8,089 cells). Venn diagram (left): 3 SIV+ cells by 10X, 39 by Catch*ReSeq, 0 shared. Counts ln(x+1)-transformed and rescaled in arbitrary units (red = 1). Contour fill (red) is the number of SIV+ cells per 1% of the UMAP bounding-box area. K. Correlation of SIV expression in single cells from (J) between 10X and Catch*ReSeq (n = 8,089 cells; Pearson R^2^ = 0.00; no cells positive in both methods). Marginal histograms on a pseudo-log_10_ scale.

To benchmark Catch*ReSeq’s sensitivity and specificity against 10X, we applied it to increasingly challenging detection tasks. First, we targeted a macaque gene, *KLRB1*, that was expressed in only a subset of cells (156 / 8614 cells) and spanned a range of expression levels in Rhesus macaque CD4^+^ T cells. Catch*ReSeq accurately recapitulated the single-cell *KLRB1* expression profile from sequencing the 10X library from the same sample at the recommended depth (333 / 345 or 97% of *KLRB1* UMIs and 145 / 156 or 93% of *KLRB1*-expressing cells), while also finding modest expression in certain cells missed by the conventional scRNAseq workflow (**Fig 1B,C**). To determine whether the incomplete *KLRB1* overlap between the two methods was stochastic or due to systematic bias, we sequenced the 10X library at over twice the recommended coverage (>40,000 reads per cell) then subsampled reads while holding Catch*ReSeq depth fixed. As 10X depth increased, cells previously detected only by Catch*ReSeq were increasingly detected by 10X as well (**Supp Fig 1C**). 78% of *KLRB1*+ cells newly detected by 10X at double the sequencing depth were already detected by Catch*ReSeq (31 / 40), and 40% of *KLRB1*+ cells detected only by Catch*ReSeq when 10X was sequenced to the recommended depth were detected by 10X when sequenced twice as deeply (31 / 77). 95% of all *KLRB1+* cells found only by one method carried only 1 *KLRB1* UMI per cell when libraries were sequenced to saturation (**Fig 1C,D, Supp Fig 1D**). These results suggest that Catch*ReSeq and 10X were subsampled from the same set of UMIs, consistent with multiple copies per UMI available for independent sampling after cDNA amplification. Together, these comparisons establish the sensitivity and specificity of the Catch*ReSeq experimental and computational pipeline.

Next, we wondered how well Catch*ReSeq could detect rare isoforms. During infection with SIV, viral RNA (vRNA) is typically less than 1% of total cellular mRNA, with individual alternatively spliced isoforms reaching a fraction of a percent^28,29^. We confirmed this observation by comparing viral to host UMIs across *in vivo* and *in vitro* SIV-infected 10X libraries and found viral UMIs were between 0.01 and 0.13% of total UMIs. To enrich for reads from SIV vRNA, we first optimized our primer design to avoid documented highly variable regions in SIV which may affect the binding of our primers (**Methods**, **Supp Fig 1E**). As a control, we performed Catch*ReSeq on an uninfected sample and detected no false positives (**Fig 1E, Supp Fig 1F**). We then performed Catch*ReSeq on cDNA from rhesus macaque CD4^+^ T cells either 7 days after infection *in vitro* or 14 days after infection *in vivo* with SIVmac239M2 (**Table 1**). These samples are referred to collectively as acute infection (in contrast to ART-suppressed).

Catch*ReSeq recapitulated 99.3% of the SIV signal detected with 10X from *in vitro* infection (16,686 / 16,808 UMIs) while revealing 56% more SIV transcribing cells (283 versus 181) and 98.7% of the *in vivo* signal (32,113 / 32,423 UMIs) with 89% more transcribing cells (1,044 versus 551 cells) (**Fig 1F,H**, **Supp Fig 1F,G**). Viral signal detected by the two methods showed a strong correlation (*in vitro*: R^2^ = 0.70 all, 0.92 non-zero values; *in vivo*: R^2^ = 0.59 all, 0.87 non-zero values) (**Fig 1G,I**), and Catch*ReSeq SIV counts per cell were uncorrelated with total RNA counts per cell (**Supp Fig 1H**).

We next asked how well Catch*ReSeq could perform on rare targets in rare cell populations. On ART, infected cells range from about 10-10,000 per million cells across individuals^30^. In CD4^+^ T cells from an ART-suppressed macaque after 154 days of treatment (**Table 1**), 10X alone detected only three virus-transcribing cells, too few to support any analysis of host cell state, whereas Catch*ReSeq detected 39 cells (13-fold over 10X) (**Fig 1J,K**). Interestingly, these SIV+ cells were transcriptionally similar to each other, suggesting a host cell state permissive of viral transcription during ART (**Supp Fig 1I**). The infected cells detected by each technique were mutually exclusive, which would be expected given the detection frequency (expected overlap 0.02 cells, P(no overlap) = 0.98; **Methods**). ART does not halt viral transcription in infected cells, but blocks to viral gene expression often prevent productive infection including generation of polyadenylated viral transcripts^13^. Nevertheless, we recovered vRNA in dozens of cells without custom capture of non-polyadenylated transcripts. This enhanced yield was specific to Catch*ReSeq as alternative approaches using viral-specific spike-in primers at reverse transcription did not improve detection in ART-suppressed samples (**Supplementary Note 1**).

### Original isoforms are broken during capture

For HIV/SIV, spliced isoforms overlap in genomic coordinates (**Fig 2A**). 10X short-read sequencing reads aligned in these overlap regions could not be uniquely mapped to a transcript isoform (**Fig 2B**). Alternatively, reads that contain splicing events (i.e. whose alignment is split across a splice junction) can be used to map isoforms^31^, but few reads from 10X generally satisfy this criterion (e.g., 18 / 50,957 viral UMIs from *in vitro*) (**Fig 2C**), despite being sequenced to saturation (63.4% for viral reads). This *in vitro* sample was undergoing productive infection, yet splicing at donor D4, which is required for productive infection^16,32^, as well as at other major canonical junctions (e.g., D2, D3, A4, A5), were not detected by 10X (**Fig 2C**).

**Figure 2:**
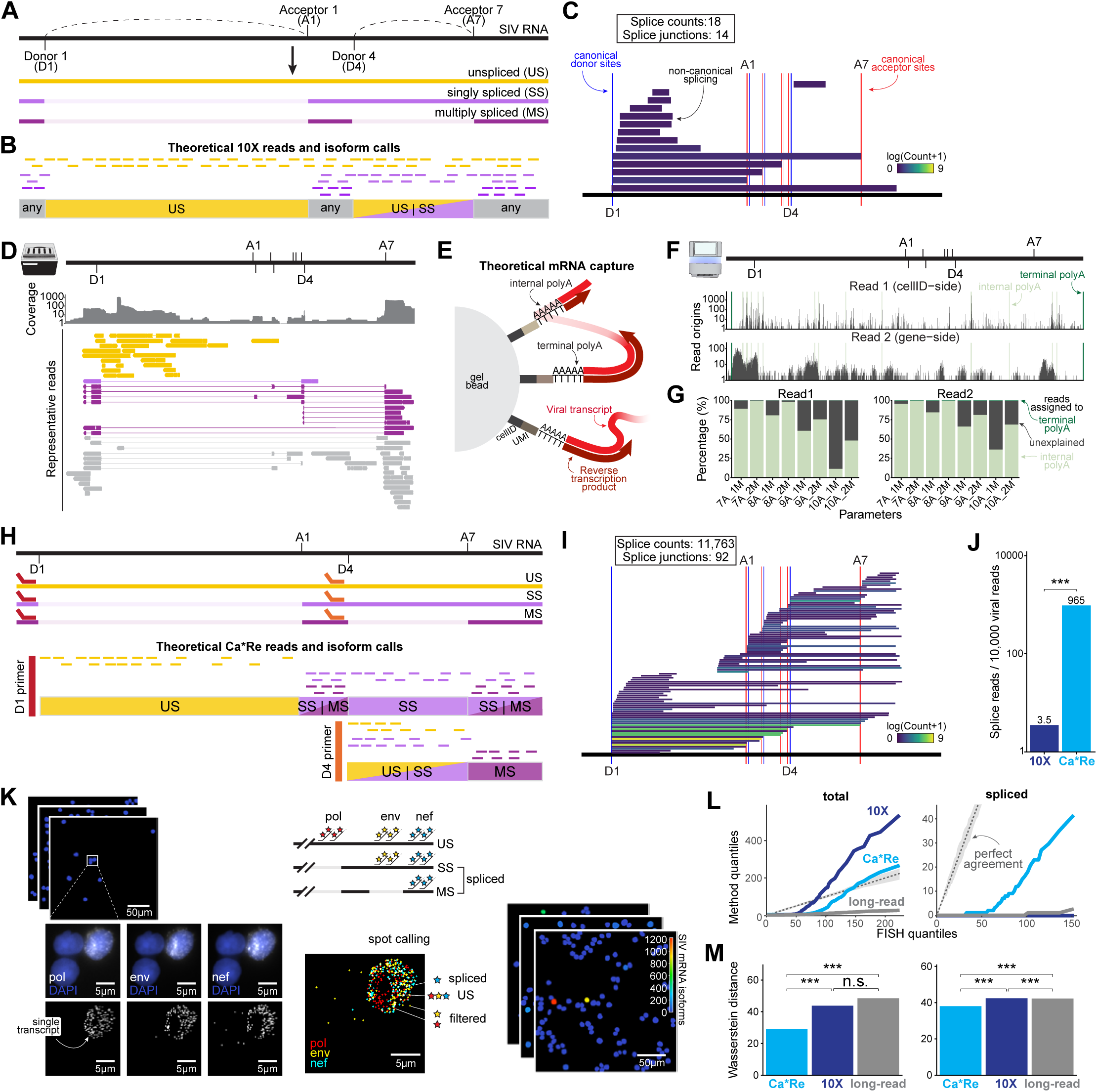
Transcript isoforms are corrupted by single cell RNAseq capture. A. Schematic representation of the major splice donor and acceptor sites used to generate the unspliced, singly spliced, and multiply spliced isoforms from full length SIV RNA. B. Schematic representation of reads from each of these isoforms after fragmentation and sequencing (top). Reads from different isoforms occupy overlapping regions, which prevents unambiguous isoform calling from 10X data (bottom). C. Reads containing a splice event mapped to the viral genome from short-read 10X sequencing (*in vitro*). Spliced reads are shown as rectangles spanning the spliced-out region, colored by log(count+1) usage frequency; canonical donor (blue) and acceptor (red) sites marked. Deduplicated and valid counts: 18 splice reads, 14 splice junctions. D. Long read sequencing (Oxford Nanopore) of *in vitro* cDNA. Top: read coverage silhouette in log-scale. Middle: representative reads colored by isoform if unambiguous, else depicted in gray. E. Schematic of theorized on-bead oligo-dT capture of mRNA molecules where both terminal and internal polyA sequences may bind to oligo-dT and prime reverse transcription. F. Short-read sequencing (Illumina) of *in vitro* infection 10X library sequenced 150 basepairs into each end. Read start location histogram of Read 1 (cellID-side) and Read 2 (gene-side) in log-scale plotted with deduplicated and valid reads, UMIs with > 1 read. Putative polyA sites 8A1M (n=16) indicated in light green, terminal polyA sites indicated in dark green in both 5’ and 3’ LTRs. G. Percentage of deduplicated reads from (F) attributed to internal polyA, terminal polyA, or of undetermined origin across internal polyA definitions. A = minimum number of consecutive A bases. M = maximum number of non-A bases i.e. mismatch bases. Reads were assigned to a polyA site if they were within 500 bases upstream for Read2 or 100 bases upstream for Read1. H. Schematic representation of the Catch*ReSeq primer strategy (top). After fragmentation and sequencing, reads from each of these primers (middle) occupy semi-distinct regions based on the isoform of origin, which allows improved unambiguous isoform calling compared to 10X data (bottom). I. As in (C) for Catch*ReSeq short-read sequencing of *in vitro*: 11763 splice reads, 92 splice junctions. J. Frequency of deduplicated reads containing a splice event out of 10,000 viral reads in *in vitro* from 10X versus Catch*ReSeq: 3.5 versus 965 (18 versus 11,763 splice events). Two-proportion test of splice events / total viral molecules, p < 0.001. K. Schematic representation of fluorescence in situ hybridization (FISH) quantification of viral isoforms in a separate replicate of *in vitro* infected macaque cells. Probe sets bind to the variably included regions between splice sites (*pol* and *env*) and a constant region (*nef*). Puncta (“spots”) are deconvolved and quantified in each channel, then colocalized to call isoforms. n = 272 cells. L. Quantile–quantile plots of per-cell isoform counts (US, total spliced) from Catch*ReSeq, 10X, and long-read (panel D) versus FISH. All three sequencing based methods derive from the same *in vitro* sample, and FISH was performed on a separate replicate. Dashed diagonal = perfect agreement with FISH; gray band = 95% bootstrap confidence interval (n = 1,000 bootstraps). n cells: Catch*ReSeq = 934, 10X = 627, long-read = 339, FISH = 272. M. Wasserstein distance from FISH (integrated area between the Q–Q curve in (L) and the diagonal), with smaller values being closer to FISH. Within-bar significance: permutation test of distribution difference from FISH (H_0_: identical). Brackets: paired-distance permutation test comparing methods’ proximity to FISH (H_0_: equally distant). *p<0.05, **p<0.01, ***p<0.001, n.s. = not significant; n = 2,000 permutations.

To resolve these disparate findings, we asked if long-read sequencing of 10X cDNA, which is often deployed for isoform calling given its longer sequence context^21,33–35^, could recover SIV isoforms. Long-read sequencing of 10X cDNA from *in vitro* infection confirmed the presence of spliced SIV transcripts (**Fig 2D**). However, we observed numerous short, apparently “fractured” reads (maximum viral read alignment 1,799 bp, median 483 bp), including reads terminating far from the single polyA site (**Fig 2D**), evidence that at least some of these molecules are not byproducts of incomplete reverse transcription. Consequently, only 54% of molecules from long-read sequencing could be confidently mapped to a single isoform (150/280), of which 77% mapped to the unspliced isoform (116/150). These results suggest that the original mRNA template gives rise to fractured cDNA at some step during cDNA generation, and demonstrate the consequences of fractured cDNA molecules for viral spliceform reconstruction.

Adenine-rich stretches can hybridize with the oligo(dT) capture sequence during mRNA capture^36–38^, placing the UMI and cell barcode at these internal polyA sequences and dividing the original transcript (**Fig 2E**). HIV and SIV are particularly rich in adenines^39^ (e.g., 34.0% SIVmac239 versus 26.0% for the average rhesus macaque protein-coding gene; 35.1% HIV-1 HXB2 versus 25.6% for the average human protein coding gene) (**Supp Fig 2A**), leading us to question if internal polyA capture could explain both the fractured molecules observed in our long-read sequencing and the broad read coverage across the viral genome in our short-read sequencing (**Fig 2F, bottom**). To test this, we sequenced a 10X library from a viremic *in vivo* samples 150 basepairs into the 3’ end, past the UMI and cell barcode into the oligo(dT)-captured sequence (**Fig 2F**). We scanned three parameters to define internal polyA sites: minimum number of consecutive adenines (A), maximum number of non-adenine bases (M), distance of read end from polyA (D) (**Methods**). Across every definition, internal A-rich regions in aggregate were captured 80 times more than the terminal polyA tail in SIV transcripts and over 6 times more for host transcripts (**Fig 2G, Supp Fig 2B,C**), while long-read data showed 5 times more viral molecules ending at internal than terminal polyA (**Supp Fig 2D**).

Internal polyA capture in SIV exceeded randomly positioned capture at every parameter set tested (**Supplementary Note 2**). These data suggest that A-rich regions in transcripts across the host and especially SIV transcriptomes were preferentially captured by oligo(dT), leading to breakage of the original isoform. Consequently, the original isoform is unrecoverable from conventional short-read sequencing and only partially recoverable from long-read sequencing.

### Catch*ReSeq captures alternative splicing of overlapping exons

Although fractured, we found evidence of spliced molecules in long-read sequencing of scRNAseq cDNA (**Fig 2D**). We asked if Catch*ReSeq could recover isoform resolution from rare alternatively spliced transcripts to create short-read libraries with viral isoform resolution. To this end, we formulated two complementary strategies: “exon-targeting,” where the variably included exon is targeted for enrichment to quantify its inclusion in transcripts, and “splice-targeting,” where the splice site immediately upstream of that exon is targeted for enrichment to quantify splicing in and out of the variably included exon. The splice-targeting strategy leverages the high rates of internal polyA capture and the relatively large distance between splice donor and acceptor sites that result in cDNA molecules terminating within the transcript body. The original isoform of a Catch*ReSeq enrichment product is assigned based on its read alignment: a molecule that used the targeted splice site would result in reads downstream of the splice acceptor (for example, a multiply spliced molecule enriched at D4 would align downstream of A7), while a molecule that did not use the targeted splice site would align within the variably included region (for example, a singly spliced molecule enriched at D4 would align upstream of A7) (**Fig 2H**, **Supp Fig 2E**). We also used reads containing splice events to call isoforms.

Catch*ReSeq recovered significantly more viral reads with splice events normalized to viral reads than 10X (965 versus 3.5 per 10,000 viral reads), including canonical splice junctions not detected in 10X (e.g. D4–A7) (**Fig 2I,J**). We observed splicing with both methods at non-canonical sites, especially located between D1 and A1, a splicing dynamic previously only observed from bulk methods^40^. We found little evidence to suggest that the non-canonical splicing we observed derived from defective provirus^41^ (**Supplementary Note 3**, **Methods**). In total, Catch*ReSeq successfully recovered both singly spliced and multiply spliced SIV isoforms from short-read sequencing (3,921 / 29,796 viral reads passing QC) whereas 10X was largely unable to resolve any spliced isoforms (8 / 50,924).

Spliced isoform products Tat and Rev are required for the positive feedback of viral transcription that supports productive infection. Consistent with this, the spliced fraction of viral UMIs increased with viral transcript counts per cell (i.e., viral transcript load) *in vitro* (Spearman rho = 0.35, p < 0.0001), and the trend strengthened among UMIs of matched read depth (9–20 reads) (Spearman rho = 0.50, p < 0.0001), arguing against sequencing depth as the driver. Correspondingly, cells with spliced viral isoforms were more frequent at high viral transcript load *in vivo* (**Supp Fig 2I**).

We next asked whether this relationship was strong enough for high viral transcript load in 10X to substitute for Catch*ReSeq spliced isoform detection. The viral expression distributions between cells with and without spliced isoforms overlap, with median recall for spliced isoform-containing cells achieved at any SIV expression cutoff tested in any sample was 0.58 (*in vitro* = 0.60; *in vivo* = 0.75 and 0.56; ART = 0; average precision *in vitro* = 0.80, *in vivo* = 0.76 and 0.56) (**Supp Fig 2F,G**). No feature of the 10X expression distribution indicated where the optimal cutoff from the precision-recall curves lay (**Supp Fig 2G**), and recovering spliced isoform-containing cells by expression threshold drew in several false positives from highly-viral-expressing cells without spliced isoforms (**Supp Fig 2H**). Notably, thresholding failed completely under ART (recall = 0). SIV expression alone therefore cannot recapitulate the Catch*ReSeq spliced isoform designation, demonstrating the unique ability of Catch*ReSeq to identify viral splicing programs within infected cells.

### RNA FISH confirms viral spliced isoform content in infected cells

To provide an orthogonal basis for viral transcript abundance and distribution, we turned to single-molecule fluorescence in situ hybridization (FISH), considered the gold standard for absolute RNA quantification and validation of RNA sequencing validation. We used limited amplification Hybridized Chain Reaction (HCR)^42^ with custom probes designed for two regions removed by SIV splicing (“pol”, “env”) and a constitutive region (“nef”)^12^, resolving each transcript as unspliced, singly spliced, or multiply spliced by probe colocalization and enabling absolute quantification of the major SIV isoforms in single cells from *in vitro* infection (**Fig 2K**, **Methods**). We compared isoform capture across three sequencing-based methods (Catch*ReSeq, 10X, and long-read) from a single *in vitro* sample to FISH from two independent *in vitro* infection replicates. For spliced and unspliced isoforms, Catch*ReSeq outperformed both 10X and long-read in count similarity to FISH as quantified by Wasserstein distance for total counts and total spliced isoforms between FISH and each method (**Fig 2L,M**). Catch*ReSeq also performed comparatively well with granular classification within spliced isoforms (singly and multiply spliced) (**Supp Fig 2J**).

For downstream analyses, we used a binary classification of isoforms in cells: infected host cells with spliced isoforms (singly or multiply spliced) detected are referred to as splice+, and cells without spliced isoforms detected (unspliced or ambiguous only) are referred to as splice-. We combined all SIV+ cells across 10X and Catch*ReSeq within each sample for downstream analyses.

### Host state and viral splicing vary along a viral transcript load continuum

HIV/SIV infected cells constitute a heterogeneous population^43^ with no unifying biomarker, although certain cell features are enriched in infected cells^17,44^, namely activation and exhaustion markers. Indeed, we detected SIV+ and splice+ cells broadly across CD4^+^ T cell transcriptional space, with certain regions enriched for viral expression and splicing (**Fig 1F,H,J**; **Fig 3A,B,G**). As different T cell subsets have varying susceptibility to infection^45^, we used consensus non-negative matrix factorization^46^ (**Methods**) to identify gene expression programs and test their enrichment between infected versus uninfected cells and between splice+ versus splice-cells (**Fig 3A–D**). *In vitro*, infected cells were enriched in a resting/memory CD4^+^ T cell gene expression program (Pattern 1; padj < 0.0001) (**Fig 3C**) marked by TCR signaling and costimulatory genes (*SOS1*, *RIPOR2*, *PTPRC* (CD45), *TAB2*, *ICOS*) and stem-like factors (*LEF1*, *SATB1*, *BACH2*), several of which are documented HIV/SIV interactors or targets of viral upregulation^47–51^. *In vivo*, infected cells were enriched in an activation/adhesion CD4^+^ T cell program (Pattern 2; padj < 0.0001) (**Fig 3D**) marked by adhesion and co-stimulatory genes (*ITGAL*, *ICOS*, *TNFRSF9*), by negative regulators of signalling (*ARHGAP15*, *RAP1GAP2*, *NEDD4L*), and by enrichment of a cytotoxic EOMES/TOX program (Pattern 4; padj = 0.0020).

**Figure 3:**
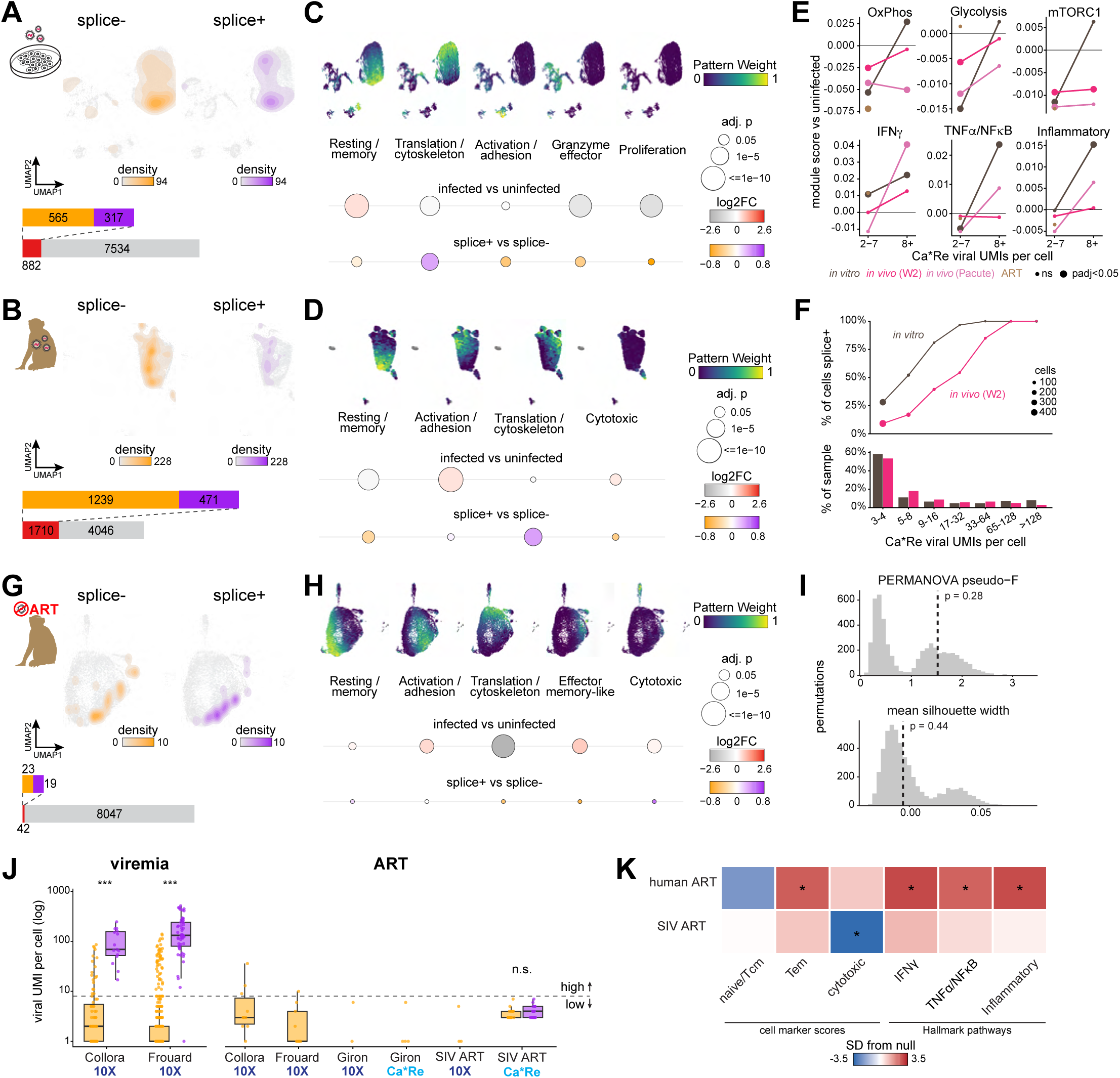
Cell states associated with differential viral isoform expression. A,B. UMAP from *in vitro* (A) and *in vivo* (n = 1, W2) (B) infection, highlighting cells with detected SIV transcripts (“infected”, left), and among those, cells with versus without detected spliced SIV transcripts (“splice+” versus “splice-”, right). A: 8,982 cells, 353 SIV+ (67 splice+, 286 splice-). B: 6,379 cells, 615 SIV+ (40 splice+, 575 splice-). C,D. Gene expression programs calculated by cNMF on highly variable genes from *in vitro* (C) and *in vivo* (D) infection. Data were first filtered to remove cells that had highest pattern weight in a non-T cell pattern, then cNMF was run again on filtered data. Annotations based on top genes with highest z-score and GSEA pathways. Below, pattern enrichment for infected versus uninfected subset (top) and splice+ versus splice-subset (bottom). *p < 0.05, **p < 0.01, ***p < 0.001, n.s. = not significant. E. Hallmark module scores of SIV+ CD4^+^ T cells detected by Catch*ReSeq, split by viral load i.e. viral UMIs per cell (2–7 or ≥8). Each point is that group’s mean score minus the mean of the uninfected CD4^+^ T cells in the same sample. Large points mark BH-adjusted Wilcoxon rank-sum p < 0.05 versus uninfected cells; groups with fewer than 10 cells are not shown. F. Viral splicing detection against per-cell viral burden for *in vitro* and *in vivo*. Top: fraction of Catch*ReSeq-detected infected CD4^+^ T cells called splice+, binned by viral UMIs per cell; point area is cells per bin, bins under 10 cells omitted. Bottom: distribution of the same cells across those bins. G. UMAP from SIV-infected ART-controlled cells highlighting cells in which SIV transcripts were detected (”infected”, left). Of those, which cells have or do not have spliced SIV transcripts detected (”splice+” versus “splice-”, right). H. Gene expression programs calculated by cNMF on highly variable genes from ART-suppression. Data were first filtered to remove cells that had highest pattern weight in a non-T cell pattern, then cNMF was run again on filtered data. Annotations based on top genes with highest z-score and GSEA pathways. Below, pattern enrichment for infected versus uninfected subset (top) and splice+ versus splice-subset (bottom). *p < 0.05, **p < 0.01, ***p < 0.001, n.s. = not significant. I. Separation of Catch*ReSeq-called CD4^+^ T cells from the ART-suppression (n = 39; 19 splice+, 20 splice-) in top 30 principal component space. Separation was quantified by PERMANOVA pseudo-F and mean silhouette width given by dashed line, tested against splice label shuffled null (5,000 permutations, gray). J. Viral UMIs per cell for every vRNA+ CD4^+^ T cell in viremic donors (left) and ART-suppressed donors (right), split by splice status (splice+, purple; splice-, orange). Columns give the dataset and the detection chemistry (10X, dark blue; Catch*ReSeq, blue). Boxes (where n ≥ 5) show the median and interquartile range, points are individual cells. Dashed line at 8 viral UMIs is the low/high viral transcript load boundary. Significance is a two-sided Wilcoxon rank-sum test of viral UMIs between splice+ and splice− cells: *p < 0.05, **p < 0.01, ***p < 0.001, n.s. = not significant. K. Host state of vRNA+ CD4^+^ T cells on ART-suppression across three CD4^+^ T cell subset marker scores and three Hallmark pathway scores for pooled human ART samples (Giron, Collora, Frouard) and SIV ART sample. Scores are percentiles within each cell’s donor/library. Tile fill is the mean RNA+ percentile expressed in SD of a null of 10,000 matched draws of equal number of RNA-cells, color limits ±3.5 SD. Asterisks: two-sided permutation p < 0.05 after Benjamini–Hochberg correction across the 15 features in that row.

Both trends replicated in a second viremic animal (**Supp Fig 3A**) and all enrichments were unchanged against uninfected cells matched on host library size (**Supplementary Note 4**, **Supp Fig 3B**).

Because the viral reservoir contains cells spanning a wide range of viral transcriptional activity^13,52^, we asked whether host state varied with viral transcript load. In low viral transcript load cells (< 8 viral UMIs) from acute samples, 72% of the ∼160 oxidative phosphorylation genes were coordinately downregulated (mean 5–10% per gene) against an expression-matched random-gene baseline (FDR 0.004) (**Fig 3E**). Glycolysis (∼90 genes), mTORC1 signaling (∼130 genes), and interferon-γ response (∼125 genes) were similarly reduced in several samples. In contrast, high viral transcript load cells (≥ 8 viral UMIs) showed increased response to infection *in vitro* through a core set of 7 genes in the inflammatory response and TNFα signaling via NFκB pathways (*IL7R*, *PTGER4*, *TNFSF10*, *CD69*, *BCL2A1*, *SLAMF1*, ZFP36), all of which were upregulated in high viral transcript load cells relative to both low viral transcript load and uninfected cells (padj < 0.01 and |log2FC| > 0.5) (**Fig 3E**). Metabolic gene suppression persisted at high viral transcript load *in vivo* (56% and 64% of oxidative phosphorylation genes down, FDR 0.007) but was reversed *in vitro* (36% down; module score +0.027, padj = 0.00076).

Which cells are called “infected” determines which host states appear associated with infection. Restricting the analysis to 10X-detected cells associated infected cells with an effector-differentiated state, with resting/memory program depleted and activation/adhesion and granzyme-effector programs enriched *in vivo* (all padj ≤ 0.031). These observations reproduced the infection phenotype reported by single-cell studies^22,53–55^, an effect driven by 10X’s biased detection of high viral transcript load cells (SuppFig 2C, **Supplementary Note 5**). With recovery of low viral transcript load cells, Catch*ReSeq revealed the resting/memory program was no longer depleted (viremia and *in vitro* upregulated, padj ≤ 0.00042; second viremia padj = 0.94). Furthermore, a translation/cytoskeletal program was suppressed (all padj < 0.0001), in concordance with observed negative regulation of mRNA translation in cells from PWH sorted for HIV DNA+ rather than vRNA load^27^.

### A translational host program is enriched with viral splicing beyond proviral transcription

We next assessed what host states were associated with splice status. Splice+ cells differed from splice-cells in host program usage (**Fig 3C,D**). *In vitro* and in cells from viremic animals, splice+ cells were associated more with the translation/cytoskeleton program than splice-cells (padj < 0.0001). This program was depleted in infected cells overall, so its enrichment distinguishes splicing rather than infection. In cells from viremic animals, splice-cells used the resting/memory program more than splice+ cells (padj = 0.013). This difference disappeared at matched viral transcript load (padj = 0.57 and 0.61) and reversed *in vitro* (padj = 0.021), indicating it reflected viral transcript load rather than splice status. In contrast, the translation/cytoskeleton program remained associated with splice+ status after accounting for viral transcript load (padj = 0.0028), thereby identifying a new host program that tracks splicing independently of proviral transcriptional activity. Notably, this decoupling was not possible with existing single-cell approaches, which detected predominantly high viral transcript load cells (**Supp Fig 2C**).

The proportion of infected cells with detectable viral splicing differed between the *in vivo* and *in vitro* infection conditions. Although a larger fraction of CD4^+^ T cells was infected *in vivo* than *in vitro* (15.8% versus 8.3%), a smaller fraction of infected cells was splice+ (27.7% versus 51.7%) (**Fig 3A,B**). In both settings, splice detection rose with viral transcript load, reaching 100% in cells carrying more than 64 viral UMIs (**Fig 3F, top**). *In vivo*, infected cells carried fewer viral transcripts, with only 14.2% carrying at least 32 viral UMIs, compared with 19.9% *in vitro* (**Fig 3F, bottom**). At matched viral transcript load, *in vitro* cells were splice+ more often than *in vivo* (**Supp Fig 3C**). *In vivo* infection was therefore broader but less productive per cell, which may suggest a selective loss of productively infected cells in the *in vivo* environment.

### Prevalent splicing without high viral transcription during ART suppression

A central question towards a cure for HIV is understanding the mechanisms by which cells with fully intact, integrated proviruses persist, even though some continue to produce complete and even spliced viral transcripts^56^. With 10X alone, we identified no splice+ cells on ART in our ART-suppressed SIV sample nor in publicly available datasets from PWH on ART, including one with enhanced viral-specific capture^22,54^ (**Supp Fig 3D**). With Catch*ReSeq, however, we successfully detected splice+ cells in nearly half of our virus-transcribing cells from SIV infected rhesus macaque on ART (19/42, 45%)(**Fig 3G**), indicating a widespread permissivity to splicing. All viral splice junctions detected were canonical, giving no evidence for transcription of replication-incompetent provirus (**Supplementary Note 3**). We detected no D4–A7 splicing in any cell from ART-suppression, though several reads contained D2–A3 splicing, which is uniquely present in spliced *tat* transcripts^11^.

All virus-transcribing cells recovered from ART-suppressed macaque were low viral transcript load (maximum 5.6 UMIs per 10,000 Catch*ReSeq reads per cell), in contrast to acute infection where splice+ cells were predominantly high viral transcript load (maximum 241 per 10,000 Catch*ReSeq reads per cell) (**Supp Fig 2H**). At the same low viral transcript load, cells from ART-suppressed macaque were 1.7–8.5 times more likely to be splice+ than cells taken during acute infection. To our knowledge, this is the first report of spliced viral transcripts in single-cell host-virus profiling of unstimulated ART-suppressed samples. Virus-transcribing cells on ART downregulated oxidative phosphorylation genes relative to uninfected cells matched on cluster and host library size (Normalized Enrichment Score (NES) −1.86, p ≤ 0.0001; NES −1.76, p = 0.0002), similar to the metabolic gene suppression seen in acute low viral transcript load cells (**Fig 3E**). This result is consistent with our finding that virus-transcribing cells in ART-suppressed macaques are metabolically quiescent relative to negative cells^25^. The 10X-detected cells alone showed no such shift (p = 0.50).

Splice+ and splice-cells from ART-suppressed macaque were indistinguishable by host cell state: 0 of 7,477 genes were differentially expressed, no cNMF program differed between the groups (minimum BH p = 0.84) (**Fig 3H**), and PERMANOVA pseudo-F and mean silhouette width in principal component space did not differ from the permutation null (p = 0.28, p = 0.44) (**Fig 3I**). At the same time, splice+ state detection was not driven by viral transcript load nor host depth (p = 0.17, p = 0.20). These results demonstrated prevalent viral splicing in macaque cells under ART-suppression and low viral transcription, suggesting splicing may not be the limiting step in maintaining sub-threshold viral transcription.

We asked how viral splicing and associated host states observed in SIV-infected macaques replicated in PWH. Similar to the splice analysis performed for 10X, we performed splice-aware reanalysis of published PWH datasets^22,54^ (**Methods**). Our analyses replicated the relative HIV+ cell frequencies from published analyses (**Supplementary Note 6**). In line with our SIV-infected macaque Catch*ReSeq results, splice+ and splice-cells from PWH were indistinguishable in host transcriptional state during viremia (**Supp Fig 3E**). Splice+ status in PWH was concentrated in high viral transcript load cells during viremia (**Fig 3J**) though rare low viral transcript load splice+ cells were detected, indicating spliced transcript detection is limited by sampling depth rather than viral transcript load (**Supplementary Note 7**). In ART-suppressed donors, reanalysis showed that HIV+ cells were largely low viral transcript load (21 / 25, **Fig 3J**), and no spliced transcripts were detected across all HIV+ ART-suppressed cells. We applied Catch*ReSeq to an ART-suppressed human cohort^57^ in which cells from PWH and HIV-negative individuals were hashed and pooled into a single reaction (**Table 2**). No HIV transcripts were detected in HIV-negative individuals (0 HIV+ / 2,363 CD4^+^ T cells from 2 HIV-negative individuals, versus 5 / 2,068 from 2 PWH). Every HIV+ cell was low viral transcript load and none revealed any spliced transcripts (**Supp Fig 3D,F,G**). Across all human ART-suppressed samples, HIV RNA+ cells were high in effector-memory identity (p = 0.031), interferon-γ response (p = 0.030), TNFα/NFκB signalling (p = 0.031) and inflammatory response (p = 0.030), and correspondingly low in naive/central-memory identity (p = 0.055) (**Fig 3K**), with the direction consistent across all three PWH datasets (**Supp Fig 3H**). The programs upregulated in HIV+ cells on ART were consistent in direction with SIV+ cells on ART, except the cytotoxic program (**Fig 3K**).

### Catch*ReSeq captures lineage-resolved splice status in T cells

We next asked how well Catch*ReSeq could recover isoform information beyond the HIV/SIV context covered thus far. Clonally heritable cell states have recently emerged as important drivers of cell fate outcomes in cancer, immune cell function, viral infections, and regeneration^58–64^. We sought to combine Catch*ReSeq with synthetic barcoding for single-cell lineage tracing to probe the timescales of isoform expression heritability in expanding primary T cells and its generalizability across fundamental T cell and disease relevant genes (**Fig 4A**).

**Figure 4:**
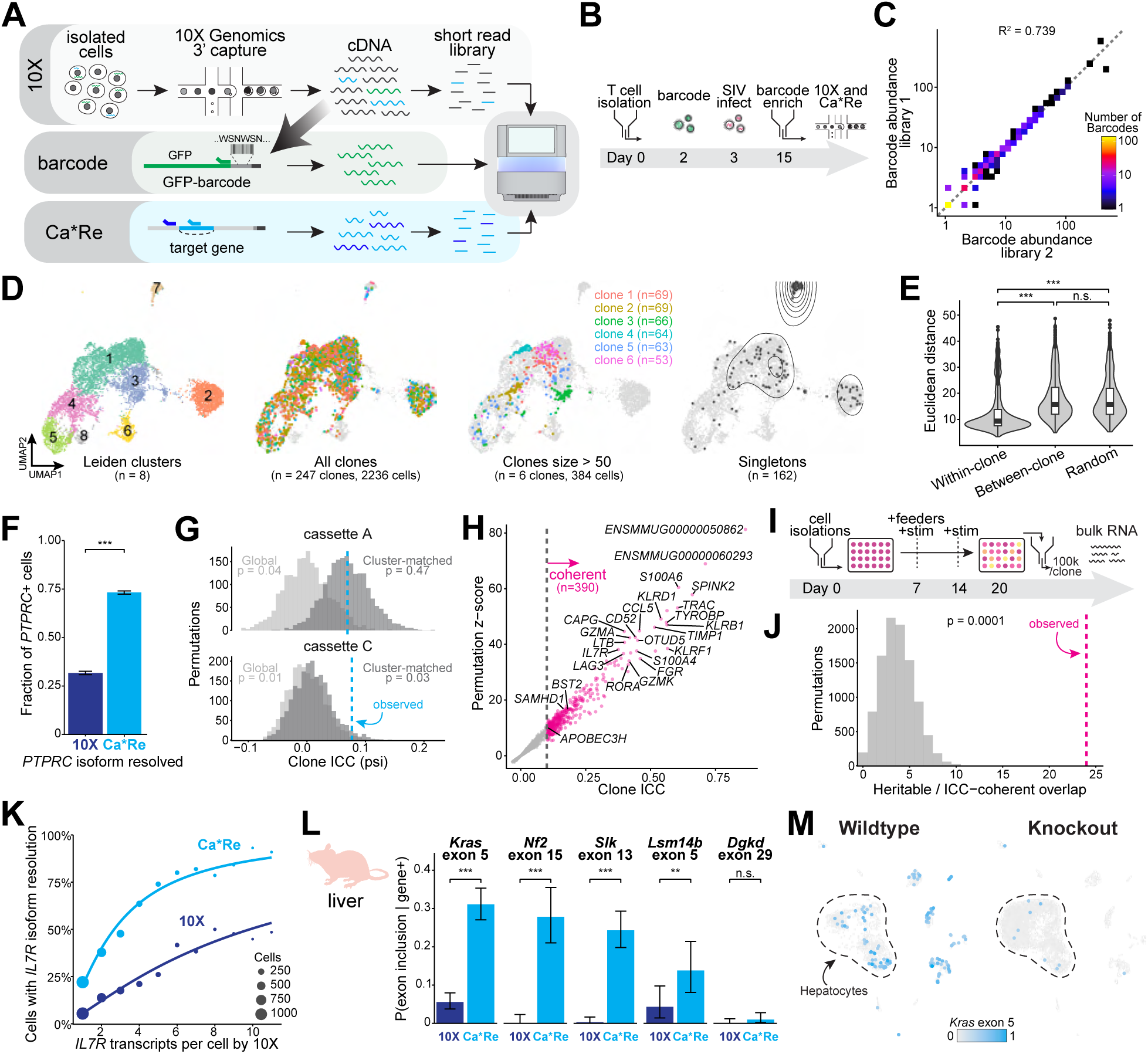
Recovering isoforms across various targets and systems. A. Schematic of Catch*ReSeq method paired with barcode recovery. B. Schematic of barcoding experiment performed on primary macaque CD4^+^ T cells. Cells were isolated from primary macaque PBMC, stimulated for two days, transduced with barcode lentivirus at low MOI, infected with SIVmac239 at low MOI, then enriched for barcoded cells by FACS before 10X. C. Abundance of each recovered barcode sequence from two independent amplifications from the same cDNA. Barcodes are binned by the number of singlet cells in which they were recovered in amplification 1 versus amplification 2 (color = barcodes per bin), with the dashed line marking equality. D. Leiden clusters, barcoded clones i.e. barcodes present in more than one cell, clones with more than 50 cells, and barcodes in only single cells (i.e. singletons), colored uniquely, imposed on UMAP. Density contours are overlaid onto singletons UMAP. E. Distribution of Euclidean distance calculated in principal component space, in order: between cells in the same clone; between cells in different clones; and between randomly sampled cells. *p < 0.05, **p < 0.01, ***p < 0.001, n.s. = not significant. F. Fraction of cells in barcoded *in vitro* sample expressing *PTPRC* gene with isoform resolved as detected by 10X via IDEIS versus Catch*ReSeq. *p < 0.05, **p < 0.01, ***p < 0.001, n.s. = not significant. G. Clonal coherence of *PTPRC* cassette splicing. Each barcoded cell in a clone of ≥2 informative cells was assigned a per-cassette psi, the fraction of its isoform-informative molecules (≥2 reads/UMI) retaining that cassette. Histograms show ICC under 2,000 clone-label shuffles, either across all clone cells (global, light grey) or restricted to cells within the same Leiden cluster (cluster-matched, dark grey). The observed value is given by the dashed line. H. Concordant gene expression between sister cells calculated by comparing the mean |log2 fold change| between sampled sister-cell pairs (x) with that of 100 random pairings of the same cells (y). Genes scored concordant (blue) showed sister-pair variation at least 10% below the shuffled control (ratio ≤ 0.9) at permutation FDR < 0.05. I. Schematic of clonal expansion experiment. See Methods. J. Heritable genes shared between clonal expansion and barcoded samples. The histogram shows the number of clonal expansion heritable genes expected to overlap the sister-concordant genes from (G) under a random null (gray, 10,000 permutations). The observed overlap is given by the dashed line. K. *IL7R* detection rate by method (10X dark blue, and Catch*ReSeq blue) versus *IL7R* gene expression (n = 2,998 *IL7R*+ cells). Points are the observed rate at each transcript count, sized by the number of cells, for counts with ≥20 cells. Logistic fit curves on log2 transcripts using all *IL7R*+ cells. L. Detection frequency by Catch*ReSeq (blue) or 10X (dark blue) of five ESRP2-spliced exons within wildtype cells expressing the parent gene. Each method scored on exon inclusion only for exon-targeting probes (*Kras*, *Lsm14b*, *Dgkd*), inclusion and exclusion for splice-targeting probes (*Nf2*, *Slk*). 10X isoform counts were quantified using BRIE2. Significance calculated using paired McNemar test. Error bars represent 95% Clopper-Pearson confidence intervals. *p < 0.05, **p < 0.01, ***p < 0.001, n.s. = not significant. M. *Kras* exon 5 scalar expression in single cells as captured by Catch*ReSeq from wildtype (left) and ESRP2 knockout (right) mouse liver in UMAP space. Counts are log_2_(x+1)-transformed and rescaled in arbitrary units. Hepatocyte clusters are encircled by the dotted line.

Across divisions from mother to daughter to large clones, cells within the same lineage can inherit states and behaviors encoded on genetic and non-genetic scales^65–67^. While TCR sequences are commonly used to label T cell clones, they are specified early in T cell development and thus lack fine temporal resolution, a limitation that synthetic barcoding addresses. Our FateMap single-cell barcoding system allowed us to track cellular responses to *in vitro* conditions and heritability of cellular state among related cells, or clones^65,68^. We barcoded macaque T cells after *ex vivo* stimulation (**Fig 4B**; **Methods**) and recovered a high proportion of original barcodes across over 2,000 barcoded clones that ranged in size from singletons, i.e., barcodes unique to a single cell, to clones of over 50 cells (**Fig 4C,D**). Cells within a clone were transcriptionally more similar to each other than cells across clones (**Fig 4E**). Barcoding was not biased to the infection status of host cells (**Supplementary Note 8**, **Methods**).

We first asked whether isoform expression of the classical T cell differentiation state marker CD45 was shared within clones or is distributed independently of clonal lineage. CD45 is coded by the *PTPRC* gene which contains three alternatively spliced exons (4, 5, and 6) whose alternative splicing and thus CD45 protein isoform expression varies across T cell differentiation state^69^. CD45 isoform expression interconverts with activation state rather than stably marking differentiation stage^70,71^, although the study of CD45 expression within clones has relied on TCR clonotype, which cannot separate antigen exposure history between clones from clone-intrinsic splicing bias. We thus asked if *PTPRC* isoform expression was shared within barcoded clones beyond what differentiation state explains. Consistent with previous reports for 3’ capture libraries, *PTPRC* isoform resolution in 10X was restricted to fewer than a third of *PTPRC*-expressing cells (3533 / 11130)^72^ (**Fig 4F**). To test whether Catch*ReSeq could provide greater sensitivity, we targeted Catch*ReSeq upstream of the three alternatively spliced exons using the splice-targeting strategy. We recovered *PTPRC* isoforms in over twice as many cells with Catch*ReSeq as 10X (8153 / 11130) (**Fig 4F**), although 75% of *PTPRC* reads did not span the entire variable region (**Supp Fig 4B**). Among isoform-resolved cells, 25% carried more than one *PTPRC* isoform (**Supp Fig 4C**). For clonally heritable expression analysis, we assigned one isoform per cell (**Methods**).

After expansion, T cell clones display variance in differentiation state^73,74^ yet carry clone-intrinsic fate biases^74,75^, but how that translates to CD45 isoform status is unknown. With barcoding, we found that *PTPRC* exon 4 (cassette A) splicing coherence within clones was fully explained by cell state (p = 0.472 against a null shuffling clone labels within transcriptional cluster), consistent with established literature^77^. In contrast, exon 6 (cassette C) splicing coherence within clones exceeded what cell state alone explains (differed from shuffled null, p = 0.032, **Fig 4G**). We further found no consistent association between *PTPRC* isoform and clone size (**Supp Fig 4D**), suggesting no correlation with expansion after *in vitro* stimulation. Together, our results show that clonal identity shapes post-stimulation *PTPRC* splicing in two distinct ways: cassette A usage tracks differentiation state, whereas cassette C usage has a dominant clone-intrinsic component. Isoform-resolved lineage tracing therefore separates splicing inherited through cell state from splicing inherited through clonal ancestry.

Heritable cell states have been implicated in host cell susceptibility to infection and resulting fates^62,63,78^. We scored each gene by the fraction of its expression variance that lies between rather than within clones (intraclass correlation, ICC; **Methods**). Clonally coherent genes included costimulatory receptors (*ICOS*, *PDCD1* (PD-1), *LAG3*, *TIGIT*), a cytotoxic/Th1 effector program (*EOMES*, *HOPX*, *KLRD1*, *KLRF1*, *TYROBP*, *IFNG*, *GZMA*, *GZMK*, *CCL5*), and antiviral restriction factors and other interferon-stimulated genes (*BST2*, *SAMHD1*, *APOBEC3H*)(**Fig 4H**). For orthogonal validation, we expanded single naive CD4^+^ T cells into clones of at least 100,000 cells and identified the most variable genes by bulk RNA sequencing (**Fig 4I**; **Methods**). Genes heritable from the clonal expansion experiment and genes concordant between sister pairs from the barcoding experiment showed significant overlap (**Fig 4J**), supporting that the Catch*ReSeq framework can be readily coupled with lineage barcoding.

We next asked whether clone specific maintenance of isoform expression were a generalized property (e.g., *PTPRC* covered above) or gene specific. We therefore tested isoform heritability for another target: *IL7R*, a gene with concordant expression across clones (**Fig 4H**) and upregulated in SIV infected cells (**Supp Fig 3A**). *IL7R* (CD127) has two protein isoforms, membrane-bound and soluble, with the transmembrane domain-containing exon 6 splicing exclusion generating the soluble form. Soluble CD127 is proposed to potentiate IL-7-driven T cell proliferation^79^ and is elevated in HIV infection, where it is suggested to be associated with disease progression^80^. We applied Catch*ReSeq to uncover which isoform was driving the concordant expression trend among clones. Catch*ReSeq recovered three times as many isoforms compared to 10X (2,664 isoforms total recovered by Catch*ReSeq versus 873 isoforms by 10X) (**Fig 4K**). Similar to *PTPRC*, soluble isoform fraction varied across *IL7R*-expressing cells (p = 0.028) but stayed a minority of each cell’s total *IL7R* transcripts (median 33% soluble). In contrast, cells appeared to express only one *IL7R* isoform at a time by 10X due to data sparsity (**Supp Fig 4E**). Too few soluble-expressing cells carried a lineage barcode to test whether the heterogeneity was clonally heritable.

### Hepatocyte subpopulations differ in sensitivity to splice regulator loss

In a final demonstration of the generalizability of Catch*ReSeq, we turned to the liver where alternative splicing by ESRP2 plays a crucial role in regeneration and homeostasis^10,81–84^. Catch*ReSeq captured more exon inclusion than 10X in four of five alternatively spliced genes in wildtype mice (4.5-to 7.0-fold, p ≤ 0.013) (**Fig 4L,M, Supp Fig 4F,G**). Alternatively spliced exons exhibited sublocalized expression patterns in wildtype hepatocytes (**Fig 4M**), which did not correlate with *Esrp2* expression (20 / 149 versus 152 / 1654 *Esrp2*+ cells, p = 0.35) and correlated mildly for host library size (R^2^ = 0.045, Spearman = 0.209, p < 0.0001). *Esrp2* was expressed as often in wildtype hepatocytes with and without Catch*ReSeq-detected alternatively spliced exons (14 / 78 (17.9%) versus 39 / 237 (16.5%)). In knockout liver, the rare cells that retained alternative spliced *Kras* exon 5 were unequally distributed, enriched in cluster 6 and absent in cluster 1 (**Supp Fig 4H–J**). This differential sensitivity to ESRP2 knockout was not driven by residual ESRP2 expression as confirmed by *Esrp2* expression knockout in these cells (1 *Esrp2* UMI / 7,002 knockout hepatocytes versus 186 UMIs / 1,803 wildtype hepatocytes; Fisher’s exact p < 0.0001). These results suggest residual splicing activity independent of ESRP2, likely resulting from adaptive remodeling of other splicing regulators^81,85,86^. Cluster 1, where splicing was ablated, showed robust expression of fetal-like program genes (*Afp*, *Anpep*, *Fxyd1*, *Gjb1)*^87^, while cluster 6, where splicing persisted, was depleted of these genes. Our result is consistent with loss of ESRP2-dependent splicing permitting fetal reprogramming in hepatocytes^81^, thereby redefining, at single-cell resolution, a link previously described through bulk approaches.

Together, these results demonstrate that Catch*ReSeq can resolve splicing and associated lineage-resolved cell states of single cells *in vivo* across biological contexts.

## DISCUSSION

Here we present Catch*ReSeq, coupling targeted enrichment of rare splice isoforms with whole transcriptome profiling in single cells from existing 10X cDNA. We applied Catch*ReSeq across several biological contexts, most consequentially to detect spliced viral transcripts in unstimulated cells under suppressive ART. Prior single-cell work relied on viral transcriptional stimulation to detect viral splicing^18^, which necessarily perturbs the permissive host cell states of interest. By measuring host state and viral splicing without perturbation, Catch*ReSeq resolves splicing in the ART-suppressed reservoir without disturbing the states that permit it. Beyond viral infection, Catch*ReSeq paired with lineage barcoding resolved the source of T cell isoform choice after stimulation, and showed that a regenerative splicing state previously described in bulk^81^ is confined to a subpopulation of mouse hepatocytes.

RNA sequencing library preparation is known to have biases that skew transcript representation relative to the actual RNA composition of the sample^88–90^. Adding to this body of literature, we document extensive cDNA fracture in oligo(dT) captured libraries across the viral and host transcriptome far exceeding reported rates^37^, and impeding and distorting isoform reconstruction from both short-and long-read sequencing. Critically, A-rich sequences are disproportionately affected, including those in HIV/SIV genomes, Alu elements and SINEs^91^, and germline-specific genes^92^. Catch*ReSeq ameliorates this biased capture, as confirmed by FISH. Two limitations remain. First, the assay requires primers against a known junction and therefore detects anticipated isoforms rather than discovering novel ones. Second, it reports the presence of a splice junction rather than isoform stoichiometry; pairing Catch*ReSeq with quantitative isoform measurement such as single-molecule FISH alongside whole-transcriptome profiling^26^ would close the remaining gap between detection and quantification.

We found splicing in 45% of virus-transcribing cells from an ART-suppressed macaque, while no spliced HIV transcripts were detected in the ART-suppressed PWH cohort. This discrepancy may be technical: of 5 virus-transcribing cells recovered across the PWH cohort, only one carried more than a single viral UMI. As splice detection scales with viral transcript load (**Fig 3F**), zero or one splice+ cell is the expected outcome even if splicing were as frequent in PWH as in macaque. Greater HIV sequence diversity may further reduce primer binding, and a biological difference between HIV and SIV cannot be excluded.If splicing were similarly prevalent in PWH, it would have a consequential impact on the interpretation of bulk data reporting frequent blocks to viral transcription at elongation and completion^13^.

Detection sensitivity also determines which host states appear associated with infection. Restricted to 10X-detected cells, infected cells appeared effector-differentiated, reproducing the phenotype reported by prior single-cell studies, an artefact of biased detection of high viral transcript load cells. Recovering low viral transcript load cells removed this signal and revealed suppression of a translation/cytoskeletal program, consistent with HIV DNA+ studies that sample the reservoir independently of viral transcript load. Inferred host state therefore depends on which cells an assay can detect. The same program was associated with splice status independently of viral transcript load, indicating that host state can contribute to splicing beyond transcript abundance.

The observed heterogeneity among virus-transcribing cells may underlie the failure of pharmacological latency reversal agents to purge the latent reservoir in ART-suppressed individuals^93,94^. By combining Catch*ReSeq with lineage tracing (**Fig 4A**), determinants of latency reversal can be resolved while accounting for host and proviral diversity. Catch*ReSeq operates on existing 10X cDNA and can therefore be applied retrospectively to longitudinal timepoints, anatomical compartments, and interventional cohorts including analytical treatment interruption, where clinical material is scarce and difficult to re-sample. These results reframe a central open question in HIV and SIV infection: whether host cell state governs viral transcriptional progression, or the viral program remodels the host cell. Catch*ReSeq paired with targeted perturbation of viral splicing and candidate host regulators provides a quantitative approach to resolving that balance.

The capture of rare isoforms from existing single-cell libraries applies to any system in which a minority of cells carries a known, consequential transcript variant. Where that variant marks a cell state of interest, Catch*ReSeq makes the transcript and its host cell measurable in the same assay. More broadly, it adds an isoform axis to libraries that were sequenced without one, supplying for annotation, modeling, and virtual cell efforts a layer of resolution that gene-level representations lack.

## ACKNOWLEDGEMENTS

We thank the New Iberia Primate Research Center and Francois Villinger for handling the non-human primate studies where the samples were sourced. We thank members of the Goyal labs for helpful discussions and comments on the manuscript, especially Jaekwon Jung for copy editing. We thank the NUseq Core facility, Feinberg Information Technology, Northwestern University Information Technology, and Quest High Performance Computing Cluster at Northwestern University Feinberg School of Medicine for their services. Flow cytometry and Illumina sequencing was performed at the Center for Synthetic Biology Biofoundry Core Facility (RRID:SCR_026869), which is supported by the Northwestern University Office for Research.

We acknowledge support from the National Institute for Theory and Mathematics in Biology (NITMB) through the National Science Foundation (DMS-2235451) and the Simons Foundation (MPTMPS-00005320). EIG acknowledges support from the National Science Foundation (NRT 2021900) and the National Institute of Allergy and Infectious Disease (NIAID) of the National Institutes of Health (NIH) (F31AI189214). YG acknowledges support from Third Coast CFAR Pilot awards, Burroughs Wellcome Fund Career Awards at the Scientific Interface, and Pew-Stewart Scholars Award. YG and EM acknowledge support from the NIAID (R01AI176599). AO acknowledges support from Ryan Fellowship. M.A-M is supported by NIH grants (R01DK146326, R01AG099681, R01DK149864, R01AG092241, R01AI189353, and R01AA031197). M.A-M is also a member of the NIH-funded BEAT-HIV Martin Delaney Collaboratory to cure HIV-1 infection (1UM1Al126620). AK acknowledges support from National Institutes of Health (R01AA010154 and R01HL126845) and Muscular Dystrophy Association (MDA1072487). NB acknowledges support from the National Institutes of Health, Chemistry-Biology Interface Training Program (T32-GM070421). JMD acknowledges support from Danica Dabich Fellowship, and DD acknowledges support from Charles F. Kade Fellowship. RD acknowledges support from the Third Coast Center for AIDS Research (CFAR, P30 AI 117943). JFH acknowledges support from the NIAID (R37AI194870, R01AI176599, R01AI174877). AK and YG are Biohub Investigators, and YG is a Biohub Senior Fellow. The content is solely the responsibility of the authors and does not necessarily represent the official views of the funding sources.

## AUTHOR CONTRIBUTIONS

YG and EIG conceived and designed the study with input from EM. EIG designed and performed a majority of the computational experiments and analyses with input from YG. EM, JK, KY, and RA designed and/or performed macaque *in vivo* studies and *in vitro* infections. EIG and GP performed library preparations. EIG and AO performed clonal experiments. EIG and AD designed and performed polyA analyses. NB, DD, and AK designed and/or performed mouse studies. EM, AK, RLR, JFH, MA-M, RD contributed to troubleshooting, collaboration-building, and advancing the project. EIG and YG wrote the manuscript with input from all authors. All authors reviewed and approved the final version of the manuscript.

## USE OF GENERATIVE AI

During the preparation of this work, the authors used Anthropic’s Opus 4.8, 5, and 5.5 and Sonnet 4.6 via Claude Code to write and edit code, and via Claude Chat to brainstorm, search literature, and proofread the manuscript. The authors reviewed and edited AI-generated content as needed and take full responsibility for the content of the published article.

## DATA AND CODE AVAILABILITY

All data associated with this manuscript is available at https://www.ebi.ac.uk/biostudies/bioimages/studies/S-BIAD4354 (imaging data) and at https://drive.google.com/drive/folders/1VZqFvq4PPePjBebWIqZQCKbCONigd2-u?usp=sharing (10X counts data). All code for the analyses in this manuscript has been deposited at https://github.com/emanuellegrody/CatchReSeq. Code for isoform calling available upon request.

## METHODS

### Macaques and sample collection

A total of 6 adult female Indian origin Rhesus macaques (Macaca mulatta; Mamu A*01, B*08 and B*17 negative; 4 years old) were used for the study described in this manuscript (**Table 1**). All macaques were selected from the colonies bred and raised at the New Iberia Research Center (NIRC), University of Louisiana at Lafayette. All animal experiments were conducted following guidelines established by the Animal Welfare Act and the NIH for housing and care of laboratory animals and performed in accordance with institutional regulations after review and approval by the Institutional Animal Care and Usage Committees (IACUC) of the University of Louisiana at Lafayette (2023-003-8841; protocol 8841-02). Humane endpoint criteria were established in accordance with the approved IACUC protocol, and animals were monitored throughout the study for signs of distress or deterioration in health. None of the animals met the criteria for humane euthanasia nor required premature termination of the study.

Rhesus macaques were housed in pairs in a biocontainment facility appropriate for SIV-infected nonhuman primates. All animals were infected with 300 TCID50 of the barcoded SIVmac239M2 stock (kindly provided by Dr. Keele at Frederick National Laboratory)^95^ intravenously and ART (Tenofovir (#FT10480) [PMPA] at 20mg/ml, Emtricitabine (#NC05645) [FTC] at 40mg/ml and Dolutegravir (#BSB37519) [DTG] at 2.5mg/ml; ART procured from Biosynth in bulk) was initiated on week 3 pi.

### *In vitro* infection

Primary macaque PBMCs were thawed and seeded at 1.5-2M cells/mL in R10MAX media supplemented with 100U/mL IL-2 and 10nM retinoic acid in anti-CD3/CD28 coated plates for three days to stimulate. Spinoculation with SIVmac239M2 and 2μg/mL polybrene was performed at 1200 x g for 2 hours at 32°C. Media was changed after 24 hours and cells were maintained at 1.5-2M cells/mL in supplemented R10MAX media. Cells were sampled, live/dead viability stained, fixed and permeabilized with the BD Cytofix/Cytoperm Kit (BD Biosciences), and stained with p24 antibody (clone KC57, NIH HIV Reagent Program) to monitor infection by flow cytometry.

#### Barcoding

High diversity barcoding lentivirus was produced as previously described^65^. For barcoded samples, cells were stimulated as described above for two days then spinoculated with barcoding lentivirus plus 4μg/mL polybrene at 1200 x g for 90 minutes at 32°C. Media was changed after 8 hours into R10MAX media supplemented with 100U/mL IL-2 and 10nM retinoic acid.

#### Clonal expansion

Frozen human PBMC isolated from fresh leukopak (Stemcell) were isolated into naive CD4^+^ T cells (Stemcell #17555) and CD3-depleted cells (Stemcell #17851). CD3-depleted cells were treated with 25μg/mL mitomycin C to generate feeders. Naive CD4^+^ T cells were single-cell sorted and cultured with feeders in Immunocult-XF media (Stemcell) supplemented with 10μg/mL IL-2, 5ng/mL IL-7, and 12.5μL/mL ImmunoCult Human CD3/CD28/CD2 T Cell Activator (Stemcell). Feeders and media were replenished at Day 7, and media was replenished at Day 14. On Day 20, clones were sorted CD4^+^ (BioLegend; clone RPA-T4) and CD69-(BioLegend; clone FN50), and stored in Qiazol at -80°C. RNA from clones with ≥100,000 cells was extracted and bulk RNA-seq libraries were generated as previously described^65^.

#### Flow cytometry and sorting

Infection frequency was quantified by intracellular p24 flow cytometry. Cell were labeled with viability dye (BD Horizon™ Fixable Viability Stain 440UV), fixed and permeabilized with the BD Cytofix/Cytoperm Kit (#554714), and stained with anti-HIV-1 p24 antibody (NIH HIV Reagent Program; clone KC57). Flow cytometry data were acquired on a Cytek Aurora spectral flow cytometer. For FACS, cells were stained with a live/dead viability stain followed by surface staining with anti-CD3 (BD; clone SP34-2) and anti-CD8 (BioLegend; clone SK1) antibodies. Cells were suspended in 1X PBS with 2% FBS and 2mM EDTA pH 8. Events were gated sequentially by scatter, viability, and CD3+ CD8-expression. Cell sorting was performed on a BD FACSAria-5 laser cell sorter (Flow Cytometry Core, Robert H. Lurie Comprehensive Cancer Center) at a flow rate of 1, or on a Sony SH800 fit with four lasers (405, 488, 561, and 638 nm) and six emission filters (450/50, 525/50, 600/60, 665/30, 720/60, 785/60).

#### scRNAseq

scRNAseq libraries (**Table 1**) were generated using the 10X Chromium Controller and Chromium Next GEM Single Cell 3’ Reagent Kits v3.1 or v4 according to manufacturer’s instructions by Northwestern University NUCore. In brief, gel beads-in-emusion (GEMs) were generated targeting 10,000 cells for recovery per sample except for D13 which targeted 7,000 and Invitro which targeted 6,000. After reverse transcription, barcoded complementary DNA (cDNA) was extracted and amplified with 11 cycles, followed by fragmentation, A-tailing, ligation with adaptors, and sample index PCR with 14 cycles.

The final 10X libraries were quantified using the High Sensitivity dsDNA kit on a Qubit 2.0 Fluorometer (Thermo Fisher) and D1000 Screen Tape on a 4200 TapeStation system (Agilent).

### Catch*ReSeq chemistry

#### Primer design strategy

To detect the inclusion of certain regions (exon-targeting), primers were targeted to the exons of interest. To detect region exclusion or splice site behavior (splice-targeting), primers were targeted upstream of known splice donor sites. All primers were determined to have high specificity for the target region via BLAST. For SIV and HIV targets, viral protein variability among sequenced SIVmac or HIV-1 clade B isolates from NCBI Virus was quantified as the probability that two randomly selected isolates differ at that position and mapped, with code adapted from Li et al. 2015^96^.

Regions with per-position amino-acid diversity ≥0.1 as well as the hypervariable regions V1-4 (Los Alamos National Laboratory; https://www.hiv.lanl.gov/content/sequence/HIV/MAP/annotation.html) were excluded from primer targets. For HIV targets, frequent 5’ mutations were additionally excluded^97^. Viral primers were targeted to major splice donors D1 and D4, with additional primers targeted for *pol*, *env*, and *nef* to ensure robust enrichment across the viral genome, relying on UMIs to deduplicate molecules that may be amplified by multiple primers. Primer sequences can be found in **Table 3**.

### PCR and library preparation

PCR amplification was performed on cDNA using a forward primer designed as described above and reverse primer that binds to a 10X homology sequence that comes after the UMI and cell barcode followed by a i5 sequencing index and Illumina P5 sequencing adapter (Table 1). Products were cleaned using left sided SPRI bead selection at 0.8X. Libraries were then fragmented and adapters added according to 10X 3’ v3.1 or v4 protocol steps 3.1-3.4. Samples were then indexed with a forward primer that binds to the adapter sequence and contains a 5’ overhang with the p7 Illumina sequencing adapter preceding the i7 index. Products were cleaned using double sided SPRI bead selection at first 0.6X then 0.8X.

#### Sequencing and preprocessing

All libraries were sequenced using NextSeq 1000/2000 XLEAP 100 cycle kits and a NextSeq 2000 sequencer (Illumina), targeted at 20,000 reads per cell for 10X libraries and 10,000,000 reads per Catch*ReSeq library, unless otherwise specified. Depending on the abundance of the target, the sequencing depth of the Catch*ReSeq library may vary. Raw sequencing data were demultiplexed using BCL Convert.

### Computational and statistical methods

#### Standard scRNAseq pipeline

Raw sequencing reads from all libraries were aligned using cellranger count to the Mmul_10 reference genome concatenated with SIVmac239 for macaque samples and GRCm39 for mouse samples. QC filtering, normalization, and downstream analysis were performed using Seurat v5.5.0. SIVmac239 reads were removed before normalization. Data were filtered to retain genes present in at least 3 cells, cells with between 200 and 3500 features i.e. genes, fewer than 20000 UMIs, and fewer than 5% mitochondrial reads. Cells that were CD4^+^CD8A^+^ were also removed.

Variable features were identified using a variance stabilizing transform for the top 2000 most variable genes. Leiden clustering and UMAP were computed from the top 30 principal components. For integrated datasets, post-QC objects were merged and re-normalized using SCTransform v2 then integrated using Harmony^98^ applied to the PCA embedding. A shared nearest neighbor graph and UMAP were computed from the top 30 Harmony components. Viral counts were zeroed in non-infectable cells, defined as CD3- and greater than one count in any of the following genes: *CD8A* (CD8+ T); *MS4A1*, *CD79A*, *CD79B*, *IGHM* (B cells); *CD14*, *LYZ*, *VCAN*, *S100A8*, *S100A9* (myeloid); *CD1C*, *FCER1A*, *CLEC4C* (DC); *PPBP*, *PF4* (platelet).

### Catch*ReSeq pipeline

A graphical summary of the computational pipeline can be found in **Supp Fig 1A**. Code is available at (https://github.com/emanuellegrody/CatchReSeq); code for Catch*ReSeq isoform calling available upon request. From the BAM files generated from alignment, UMIs and cellIDs were isolated from reads confidently mapped to regions of interest (see Isoform Calculation). Reads are then filtered to retain only cellIDs with greater than 100 UMIs in the 10X library from the same sample.

Additional filters are set to the following minimums: ≥2 [uninfected, ART] or ≥7 [*in vitro*] or ≥9 reads [*in vivo*] and ≥3 UMIs [all]. UMIs were then converted to isoform counts, log normalized, filtered for non-infectable cells, and joined to the matching 10X dataset.

### Isoform calculation

Isoform were assigned after reference genome alignment. For Catch*ReSeq exon-targeting, reads aligning to the variably included exon or downstream are retained from all samples, with libraries not targeting the variably included exon used to determine background reads. For Catch*ReSeq splice-targeting, reads spanning a splice junction (e.g. aligning upstream of donor and downstream of acceptor) were retained, while for particularly A-rich sequences e.g. HIV/SIV, reads aligning after the splice acceptor for the variably included exon and before the next expected splice acceptor are additionally retained (**Supp Fig 2E**). Viral UMIs mapping to multiple regions were called isoform unknown. For 10X, viral isoforms were assigned from read alignment (unspliced) and from junction-spanning reads, with all other isoforms calculated by BRIE2^31^.

### Sequencing saturation

Saturation was computed from the per-UMI read counts assigned to the target (*KLRB1* or SIV). Reads were rarefied by drawing Binomial(read count, f) reads at f = 0.1-1.0 in steps of 0.1, discarding UMIs with no sampled read.

Sequencing saturation was then taken as 1 − (unique UMIs / total reads) in the subsample, computed separately for 10X and Catch*ReSeq. Each depth was sampled in 5 independent replicates; curves show the mean across replicates on both axes and the band ±1 SD of saturation. For rarified detection, the same subsampling fractions and replicates were used: for Catch*ReSeq, each UMI passing min_reads at full depth was retained with probability 1 − (1 − f)(read count), and cells with at least min_umi retained UMIs inside the CD4 T gate were called positive; for 10X, reads were thinned as above and cells carrying at least two target reads inside the gate were called positive.

### Expected overlap between 10X and Catch*ReSeq

Overlap between the two methods’ SIV+ calls was evaluated against a hypergeometric null within the CD4^+^ T gate, with cells stratified into 20 equal-count bins of total RNA counts. Expected overlap is the sum over bins of (Ca*Re+ x 10X+ / cells in bin) and P(no overlap) the product of the per-bin hypergeometric P(0).

### Replication incompetence

Replication incompetence was defined as a non-canonical splicing junction in all reads in a cell aligning to the region after filtering.

### polyA analysis

The 10X library from a viremic *in vivo* sample was sequenced 150 basepairs each into Read1 (cellID-side) and Read2 (gene-side). Reads were aligned using STAR to the joint macaque and SIV reference genome described above. UMIs and cellIDs extracted from Read1 were filtered for UMI xf flag {17,25} with at least one read and cellID that survive Seurat QC as described above, respectively. For host analysis, reads were filtered for alignment to host protein coding genes. For virus analysis, UMIs with xf flag of 0 were also allowed to include reads multimapping to the 5’ and 3’LTRs.

Terminal polyA location was set at the polyadenylation signal in each LTR for SIV, the end of the transcript annotation for host genes. Internal polyA sites were defined iteratively without positional overlap by sweeping through the following parameter sets: minimum length of A’s ∈ 6-10; maximum non-A i.e. mismatched bases ∈ 0-2; and distance to putative polyA site ∈ {400, 500, 600} for Read2 and {50, 100, 150} for Read1. Reads that did not align to terminal nor internal polyA were categorized as unexplained. Parameter sets were excluded if > 50% of reads were unexplained. Separately, 20,000 positions were drawn uniformly across the SIV genome and read assignments were compared against defined internal polyA sites (**Supplementary Information**).

### Hallmark scores

Hallmark scores were expressed relative to uninfected cells after regressing out library size, number of genes detected, and ribosomal fraction.

### Gene expression programs using cNMF

Gene expression programs were determined by consensus NMF (cNMF)^46^ using 200 iterations on 2000 highly variable genes. The number of components (K) was selected to reasonably balance high stability and low error. A local density threshold of 0.1 was selected to remove outlier runs. Gene expression patterns were annotated using preranked gene set enrichment analysis (GSEA) on gene spectra Z-scores against rhesus macaque-mapped MSigDB^99^ Hallmark, Reactome, and GO:BP using fgsea (nPerm = 10,000) and msigdbr, and overrepresentation analysis on the top 50 genes by Z-score using a one-sided hypergeometric test against GO:BP with all detected genes as background.

#### Host size control

Infected and uninfected cells were binned into deciles of host library size i.e. host UMIs per cell, and within each bin up to five uninfected cells were drawn at random per infected cell in that bin. Program usage was then compared between the infected cells and the matched reference with the same two-sided Wilcoxon rank-sum test per cNMF program, BH-adjusted across programs.

### PWH data reanalysis

FASTQ files were downloaded from SRA and ENA for reanalysis: Collora et al. (BioProject PRJNA777601) unstimulated CD4^+^ T cell gene-expression libraries only; Frouard et al. (GEO GSE266455, BioProject PRJNA1107214); and Giron et al. (GEO GSE254483, BioProject PRJNA1070656). Giron et al. gene expression libraries were re-sequenced in-house. Raw sequencing reads from all libraries were aligned using cellranger count to the GRCh38 reference genome concatenated with HIV-1 HXB2. QC thresholds were set as originally published, with additional filters at ≤5% mitochondrial reads and ≤10,000 counts for Giron et al. Cells were filtered for CD4^+^ T cells by ADT data where available or by marker gene panel when unavailable. Giron et al. pools were demultiplexed to donors with HTODemux and filtered with demuxmix to remove multiplets and droplets of ambiguous assignment. CD4^+^ T cells were called HIV RNA+ if assigned at least two HXB2 reads, following original methods. Splice isoforms were called as described above for SIV 10X libraries.

### Barcode recovery and assignment

Lineage barcodes were recovered from two independently prepared barcode libraries built from the same 10X cDNA and assigned to cells as previously published^65^ (https://github.com/GoyalLab/Updated_10XBarcodeMatching). Briefly, reads were demultiplexed by sample index, filtered on stagger length and GFP primer, followed by primer base quality and homopolymer content. CellIDs were restricted to the cellranger cell barcode whitelist. Barcode sequences were collapsed with starcode^100^ (Levenshtein distance 8). Singlets were identified with singletCode^68^ and pooled across the two libraries. Cells were assigned barcodes supported by >4 UMIs, taking two-barcode combinations only where the combination recurred across cells. Three promiscuous barcodes appearing in more than three distinct combinations were removed.

### PTPRC isoform assignment

Because surface CD45 phenotyping conventionally resolves T cells into discrete isoform-defined populations, and following approaches to CD45 isoform assignment from single-cell data^72^, we assigned each cell to the dominant isoform supported by the most reads, with priority to isoforms with full information.

### Intraclass correlation

For each gene detected in at least 5% of cells belonging to a clone of at least two cells, we fit a one-way random-effects model with clone identity as the grouping factor and took the fraction of expression variance between clones rather than within them (ICC), with unequal clone sizes handled by the n₀ correction. Each gene’s ICC was compared against 100 clone-label permutations, and genes were called coherent at BH-adjusted permutation p < 0.05 together with ICC ≥ 0.10.

### Mouse liver annotation

Leiden clusters were annotated by scoring per-cluster mean log-normalized expression against a panel of canonical mouse-liver cell-type signatures (hepatocyte, periportal/pericentral hepatocyte sub-zones, cholangiocyte, LSEC, Kupffer, stellate, T/NK, B/plasma, neutrophil, erythrocyte) and assigning each cluster the top-scoring identity. Within-hepatocyte analyses were restricted to clusters annotated as hepatocyte (clusters 1, 2, 3, 4, 5, 6). Clusters with fewer than 20 target gene-expressing cells were excluded from per-cluster rate inference but pooled into the multi-axis state-score analyses.

### Clonal expansion heritable genes

Sequencing data were analysed using NextFlow rnaseq v3.17.0^101,102^ using default parameters, against the GRCh38 primary assembly. For experimental and control samples separately, transcripts per million (TPM) coefficients of variation were modeled against log_2_ TPM using Poisson GLMs. Genes with mean TPM >1.5 and residuals above the 98th percentile of experimental-sample residuals were retained.

#### Oxford Nanopore sequencing

Catch*ReSeq enrichment products before short-read library preparation underwent another 8 cycles of PCR to add -NNN overhangs for balanced strand sequencing following Lebrigand et al.^34^. Libraries were prepared using the Ligation Sequencing Kit V14 (Oxford Nanopore) and sequenced on an Oxford Nanopore GridION. Data were analysed using NextFlow scnanoseq v.1.0.0 and 1.2.1^102,103^ using default parameters, against the SIVmac239 genome.

### Fluorescence in situ hybridization

#### Probe design

HCR v3.0 probes were custom designed with Molecular Instruments for the *pol* (30 probes, lot # RAQ572), *env* (24, #RAQ573), and *nef* (9, #RAQ574) regions of SIVmac239. *Env* probes by design avoid hypervariable regions.

#### Sample preparation and imaging

The HCR protocol^42^ was modified for increased yield for suspension cells following an adapted protocol from Peterman et al.^104^. The main modifications were to use a swinging bucket rotor for all spins, reduce the number of washes and spins, and decrease deceleration on spins with probe hybridization buffer. Probe hybridization buffer, wash buffer, amplification buffer, probes, and amplifiers were purchased from Molecular Instruments HCR v3.0. Cells were fixed by incubating with 4% paraformaldehyde in PBS for 15 minutes at room temperature. Cells were next permeabilized in 0.5% Tween-20 in PBS for 15 minutes at room temperature. Cells were first primed in probe hybridization buffer for 30 minutes at 37C then incubated with custom designed probe sets in probe hybridization buffer at 4nM (e.g. 0.4uL probe in 100uL buffer) overnight at 37C. The next day, cells were spun at decreased deceleration to remove probe hybridization buffer and washed first in probe wash buffer for 15 minutes at 37C then in 5X SSCT for 5 minutes at room temperature. Cells were first primed in amplification buffer during hairpin snapcooling then incubated in the dark with 60pM (i.e. 4.5pmol in 75uL) h1 and h2 hairpins for 45-90 minutes. Cells were then washed twice in 5X SSCT at room temperature, first for 30 minutes, then 5 minutes. Finally, cells were stained with DAPI in PBS at 50ug/mL for 5 minutes, washed twice with 5X SSCT for 5 minutes each, then suspended in SlowFade Gold mountant (Thermo Fisher) in a glass-bottomed 96 well plate or slide. Samples were imaged at 60X oil immersion as previously described^65^.

### Processing and quantification

Cells were manually segmented in Fiji^105^ and segmentation masks were created with BIOP plugin. With Big-FISH^106^ (v0.6.2), diffraction-limited spots were detected and quantified from maximum intensity projections of each channel corresponding to each probe set. Detection thresholds were set per channel as the mean of automatic per-field thresholds across all fields of view in an imaging session. Spots were masked by cell segmentation. Dense spot clusters were deconvolved by intensity and transcription sites were removed. Spliced isoforms were assigned per *nef* spot based on the colocalization (within 3 spot radius distance) of spots from probe sets targeting *pol*, *env*, and *nef*:

Unspliced (US): nef AND env AND pol

Singly spliced (SS): (nef AND env) AND NOT pol

Multiply spliced (MS): nef AND NOT env

To correct misassignment due to low *env* signal, *env* intensity was measured at each MS spot as peak-over-background of background median absolute deviation, then spots exceeding a per-field threshold set from within-mask null allowing 1% false positive rate were reassigned US (if colocalized with *pol*) or SS (if not). Cells containing at least one US or SS spot were kept for downstream analysis.

