## Supplementary Information for "Catch*ReSeq resolves rare splice isoforms and their host cell states at single-cell resolution"

Supplementary Figure 1

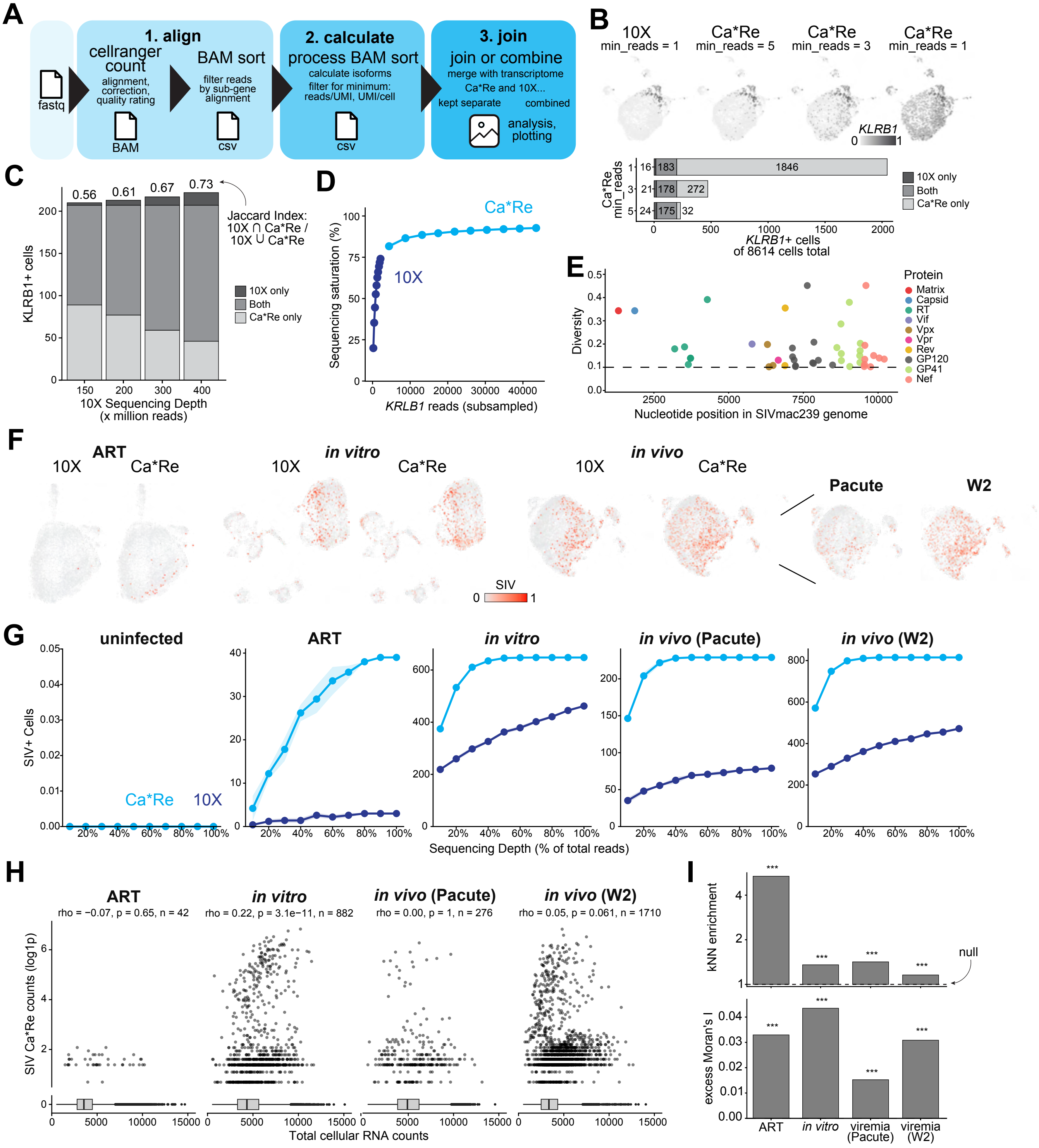

**Supplemental Figure 1: Developing the Catch*ReSeq method.**

A. Schematic of Catch*ReSeq computational pipeline.
B. Minimum reads-per-UMI threshold sweep to balance true/false positives for *KLRB1* from uninfected sample (n = 8,614 cells; 10X fixed at min_reads = 1). Top: *KLRB1*+ cells on UMAP at 10X (min_reads = 1) and Catch*ReSeq (min_reads = 5, 3, 1). Bottom: *KLRB1*+ cell counts by detection category (10X-only, both, Catch*ReSeq-only) at each Catch*ReSeq threshold.
C. *KLRB1* detection by 10X and Catch*ReSeq as a function of 10X sequencing depth (150–400 million reads) while Catch*ReSeq read depth was held fixed (min_reads = 5). Stacked bars show the number of *KLRB1*+ cells by detection category (10X-only, both, Catch*ReSeq-only). Above the bars: Jaccard index (10X ∩ Catch*ReSeq / 10X ∪ Catch*ReSeq).
D. *KLRB1* sequencing saturation (1 − unique *KLRB1* UMIs / *KLRB1* reads) versus subsampled *KLRB1* read depth for 10X (dark blue) and Catch*ReSeq (blue). Reads per UMI were thinned binomially at 10–100% of the full depth in 10% steps, 5 replicates per step; points are the mean across replicates in both axes, band = ±1 SD of saturation.
E. Per-position amino-acid (pairwise) diversity across the SIVmac239 proteome versus genomic nucleotide position, colored by protein. Diversity is the probability that two randomly chosen isolates differ at that position. SIVmac isolates sourced from NCBI Virus. Only positions with diversity ≥ 0.1 (dashed line) are plotted.
F. UMAP plots with SIV+ CD4 T cells detected by 10X (left) or Catch*ReSeq (right) plotted for each sample. The integrated viremic sample Catch*ReSeq is plotted both as one object and split by individual ("Pacute" and "W2"). Counts ln(x+1)-transformed and rescaled in arbitrary units (red = 1)
G. SIV+ CD4 T cells detected versus subsampled read depth for 10X (dark blue) and Catch*ReSeq (blue) across uninfected, ART, *in vitro*, and *in vivo* (Pacute, W2) samples. 5 replicates per depth, lines = means, band = ±1 SD. Detection thresholds 10X ≥2 reads; Catch*ReSeq ≥2 [uninfected, ART] or ≥7 [in vitro] or ≥9 reads [viremia] and ≥3 UMI [all].
H. Total cellular RNA counts versus SIV transcripts (ln(x+1)) per SIV+ cell, with SIV- cells given by box & whisker plots. Spearman correlation: ART ρ = -0.07, p = 0.65, n = 42; *in vitro* ρ = 0.22, p = $3.1\times10^{-11}$, n = 353; *in vivo* (Pacute) ρ = 0.00, p = 1, n = 276; *in vivo* (W2) ρ = 0.05, p = 0.061, n = 1710.
I. Clustering of Catch*ReSeq SIV+ CD4+ T cells on k-nearest-neighbor graphs (k = 30). Top: kNN enrichment, the mean fraction SIV+ of 30 neighbors per SIV+ cell, divided SIV+ neighbors fraction for random cell sets matched by library size. Bottom: excess Moran's I, the spatial autocorrelation of SIV+ status on the kNN graph minus that of the library size-matched random sets. Dashed line: null. *** p ≤ 0.001, one-sided, 999 permutations.

### Supplementary Figure 2

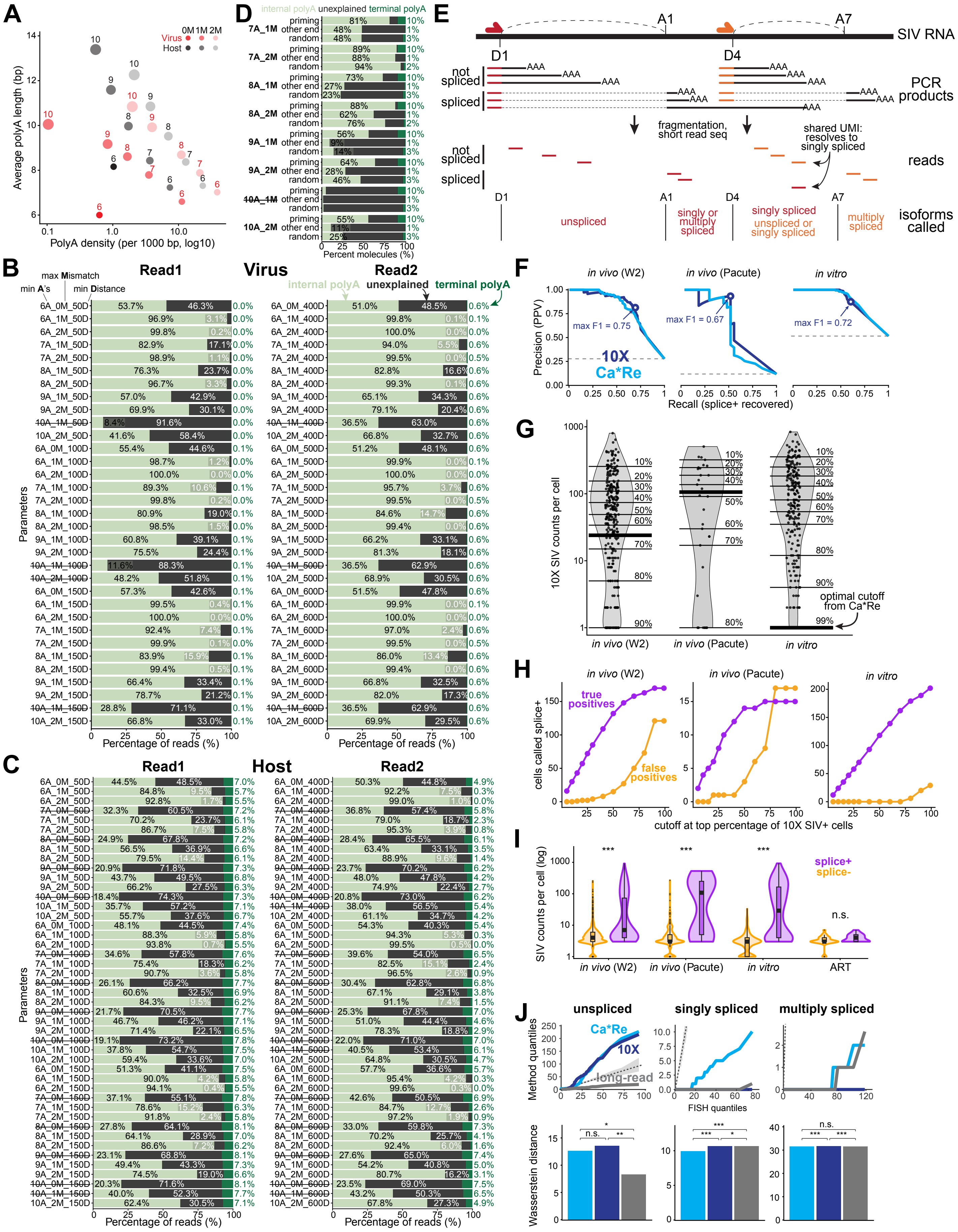

**Supplementary Figure 2: Effect of polyA capture on 10X cDNA libraries.**

A. Internal polyA tract density (tracts per 1,000 bp, $\log_{10}$ x-axis) versus average tract length (bp) for the SIVmac239 (reds) and host (grays) genomes computed from genome FASTA across parameters. Point size = minimum A-run length (6–10, labeled); transparency = maximum allowed mismatches (M = 0, 1, 2). Host genes restricted to protein-coding.
B. Percent of SIV-aligning short-reads (n = 21,165 read pairs from filtered cells, UMIs with >1 read) attributable to terminal polyA (dark green), internal polyA (light green), or unexplained (gray), across A-run length (A), mismatch (M), and distance-window (D) parameters; e.g. "8A_1M_500D" = ≥8-base A-run, ≤1 mismatch, ≤500 bp window. Terminal polyA = priming within the distance window of the canonical SIVmac239 3′ polyadenylation site. Read1 windows ≤50/100/150 bp, Read2 ≤400/500/600 bp. Strikethrough: parameter sets excluded from summary statistics for > 50% reads unexplained.
C. As in (B) for reads aligning to host protein-coding genes (n = 7.6 million reads). Terminal polyA = priming within the distance window of the 3' end of the gene annotation.
D. Percentage of SIV-aligning long-reads (n = 184 molecules) whose untemplated polyA/polyT end ("priming") lies within 100 bp of an internal polyA (light green), terminal polyA (dark green), or neither (unexplained, gray). The same classification is applied to the other ends of the same molecules and to 20,000 uniformly drawn transcript positions ("random"). Strikethrough: parameter sets excluded from summary statistics for > 50% reads unexplained.
E. Schematic of Catch*ReSeq targeting splice sites with example reads and isoform calling strategy.
F. Precision-recall curves for predicting Catch*ReSeq splice+ status from per-cell SIV expression by each method (10X, dark blue; Catch*ReSeq, blue). X axis = recall (fraction of splice+ cells recovered), y axis = precision (positive predictive value), with the dashed no-skill line at the splice+ prevalence i.e. splice+ cells / SIV+ cells. Circles mark the F1-optimal cutoff.
G. Violins of per-cell 10X SIV log counts in SIV+ cells. Light horizontal lines mark the count cutoffs that correspond to selecting the top X% of 10X SIV+ cells (5-90%). Heavy horizontal line at the Catch*ReSeq-derived F1-optimal cutoff.
H. 10X SIV+ cells above threshold scored against the Catch*ReSeq splice+ label over an SIV-expression cutoff sweep (top 5% to 100%) of 10X SIV+ cells. Purple = true positives, orange = false positives.
I. Violin plots of total SIV counts per SIV+ cell for each viremia animal, *in vitro*, and ART, with cells categorized by whether spliced isoforms were detected by Catch*ReSeq (splice+, purple) or not (splice-, orange). Counts back-transformed from log1p and shown on a log scale. Two-sided Wilcoxon rank-sum p, *p < 0.05, **p < 0.01, ***p < 0.001, n.s. = not significant
J. Top: Quantile-quantile plots comparing isoform count distribution from Catch*ReSeq, 10X, and long read (panel C) versus FISH for singly spliced and multiply spliced isoforms, which were combined for the spliced count (Fig2 L-M). Dashed diagonal represents perfect agreement with FISH. Gray shaded region shows 95% bootstrap confidence interval. Bottom: Wasserstein distance quantifying probability distribution deviation from FISH i.e. integrated area from Q-Q curve to diagonal (panel H). Smaller values indicate closer agreement to FISH. Significance within bars: permutation tests for distribution difference from FISH ($H_0$: distributions identical). Brackets above bars: paired-distance permutation tests comparing methods' relative proximity to FISH ($H_0$: methods equally distant to reference). *p < 0.05, **p < 0.01, ***p < 0.001, n.s. = not significant; n = 2,000 permutations.

Supplementary Figure 3

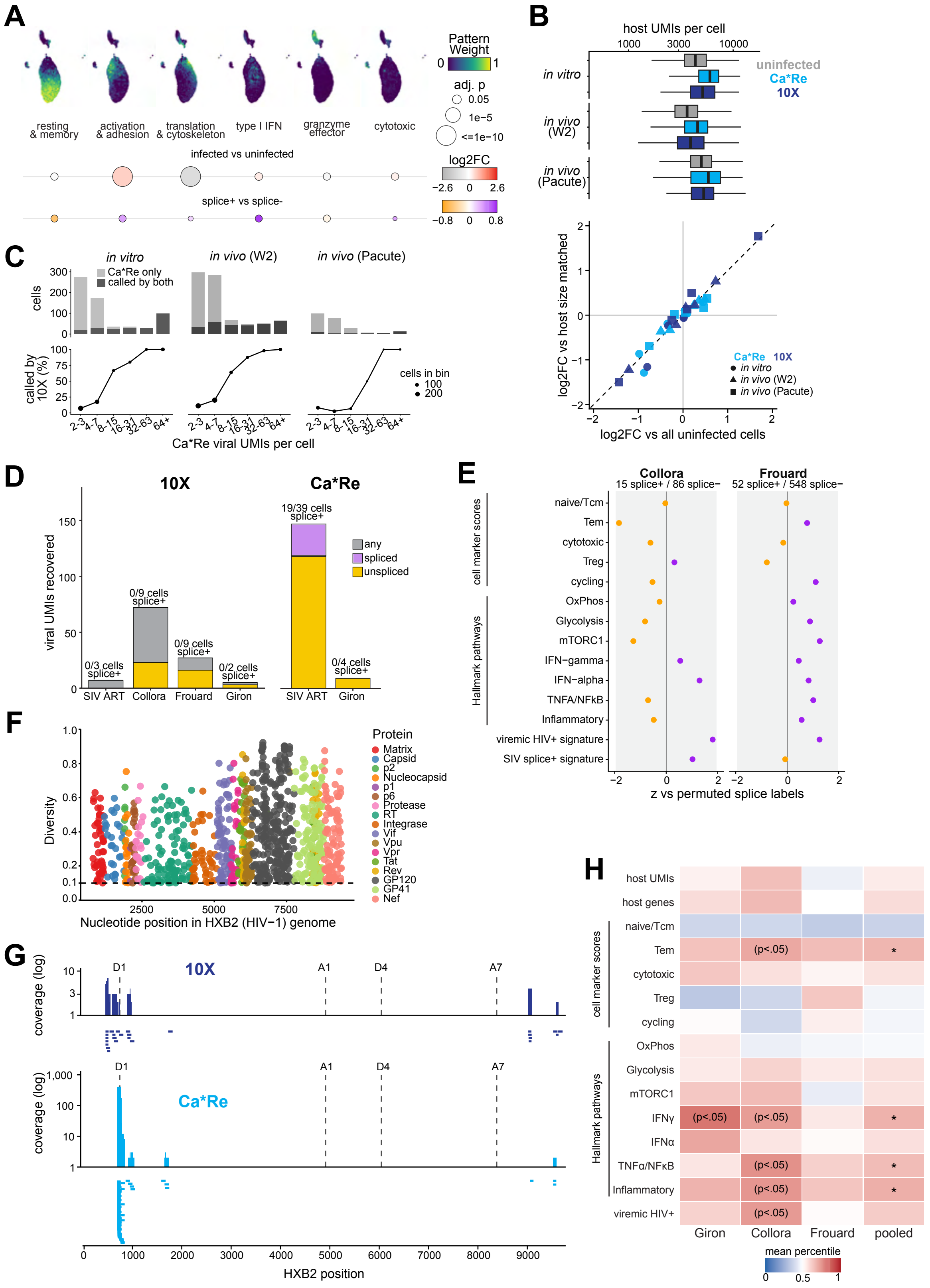

**Supplementary Figure 3: Differential gene expression analyses.**

A. Gene expression programs calculated by cNMF on highly variable genes from second *in vivo* infection animal (“Pacute”). Data were first filtered to remove cells that had highest pattern weight in a non-T cell pattern, then cNMF was run again on filtered data. Annotations based on top genes with highest z-score and GSEA pathways. Below, pattern enrichment for infected versus uninfected subset (top) and splice+ versus splice- subset (bottom). $*p < 0.05$, $**p < 0.01$, $***p < 0.001$, n.s. = not significant.
B. Top: host UMIs per cell (log scale) for uninfected CD4+ T cells, all Catch*ReSeq-positive cells and 10X-detected cells per sample. Boxes span the interquartile range with the median, whiskers to 1.5x the interquartile range, outliers not drawn. Bottom: Log2 fold change in mean cNMF program usage against uninfected cells matched on host library size decile, plotted against the same fold change computed against all uninfected cells. Matched references were drawn over ten quantile bins of host UMIs. Both axes are log2(mean usage in the infected population / mean usage in the reference). Dashed line, identity. Color gives detection method (10X, dark blue; Catch*ReSeq, blue) and shape the sample.
C. Recovery of infected cells by 10X as a function of viral transcript abundance. CD4+ T cells called infected by Catch*ReSeq, binned by Ca*Re viral UMIs per cell. Top: cells per bin, shaded by whether the 10X library prepared from the same cell suspension also called the cell (dark, called by both; light, Ca*Re only). Bottom: the percentage of each bin also called by 10X, point area proportional to the number of cells in the bin for bins with >5 cells.
D. Viral UMIs recovered from each ART-suppressed dataset, classified by isoform, detected by 10X (left) and Catch*ReSeq for the datasets that have one (right). Labels above bars give the number of called cells carrying any spliced UMI.
E. Module scores for CD4+ T-cell subsets, Hallmark pathways, a viremic HIV+ signature and an SIV splice+ signature, compared between splice+ and splice− HIV+ CD4+ T cells from viremic participants in Collora et al. and Frouard et al. Each score was adjusted for viral and host UMI counts and compared against 10,000 permutations of the splice labels within library and viral-burden bins. Points show z versus the permutation null (purple, higher in splice+; orange, higher in splice-), and the grey band marks $|z| < 1.96$.
F. Per-position amino-acid (pairwise) diversity across the HIV-1 clade B proteome versus genomic nucleotide position, colored by protein. Only positions with diversity ≥ 0.1 are plotted (dashed line). HIV-1 clade B isolates sourced from NCBI Viruses.
G. Alignment of Giron viral reads along the HIV-1 (HXB2) genome in the 10X (dark blue) and Catch*ReSeq (blue) libraries. For each method, top: All deduplicated read alignments per-position read depth on log-scale; and bottom: representative individual read alignments. Dashed lines mark the major HIV-1 splice donor and acceptor sites.
H. Host-cell state of HIV and SIV RNA+ CD4+ T cells on ART. Each feature (y-axis) was converted to a within-group percentile among the CD4+ T cells of the same donor or library ((rank − 0.5)/n), so that values are comparable across datasets of differing depth and composition. Tiles show the mean percentile of the RNA+ cells, red above the midpoint and blue below, compared against 10,000 draws of as many RNA− cells from the same donors or libraries. p is two-sided and Benjamini–Hochberg corrected across the 15 features within a column; * BH $p < 0.05$, ($p<.05$) nominal only. n = 5 (Giron), 10 (Collora), 9 (Frouard), 24 (PWH pooled), and 42 (SIV ART) RNA+ cells.

Supplementary Figure 4

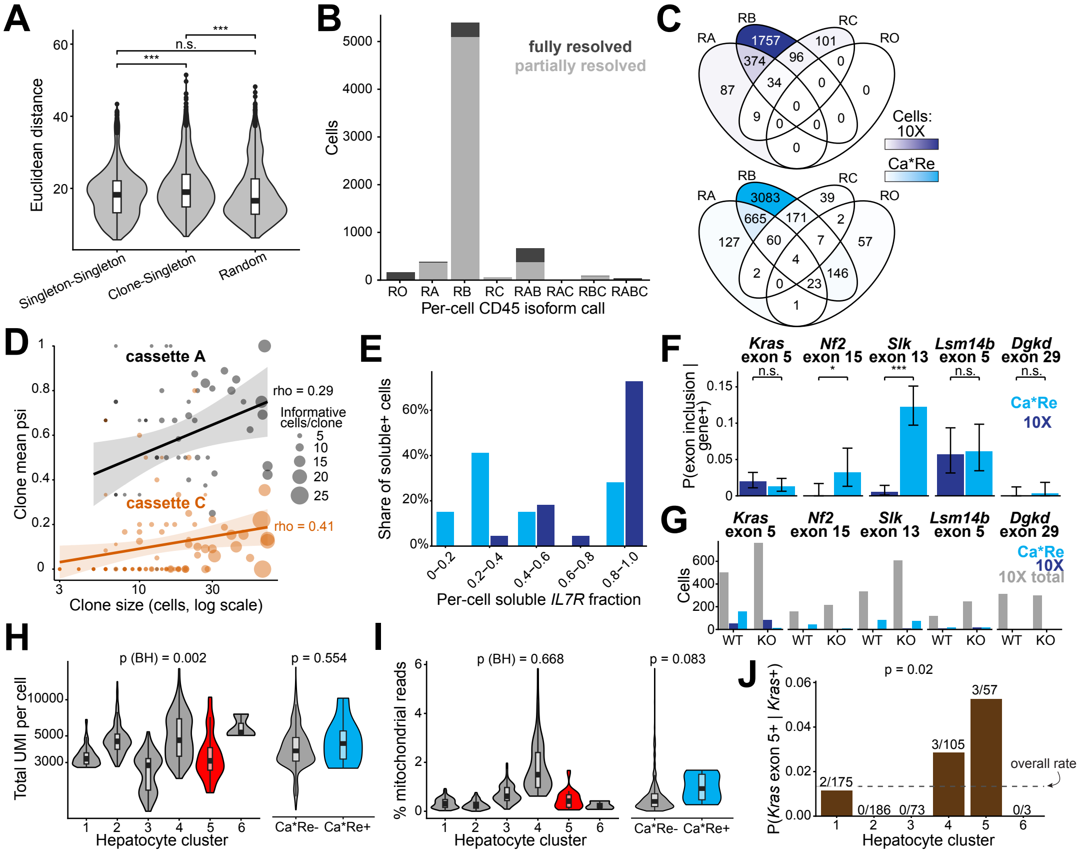

**Supplementary Figure 4: Barcode analyses and liver controls.**

A. Distribution of Euclidean distance calculated in principal component space, in order: between individual singletons; between cells in clones and singletons; and between randomly sampled cells.

B. PTPRC single-isoform reconstruction after assigning single CD45 isoform to each cell from Catch*ReSeq UMI with ≥2 reads: a molecule resolving all three variable exons wins the assignment, otherwise the most abundant isoform is used. The barplot shows the resulting per-cell isoform composition across the eight exon-A/B/C combinations, with each bar split into fully resolved (dark gray) and partially resolved (light gray).

C. Four-way Venn diagrams of per-cell *PTPRC* isoform expression using 10X counts via IDEIS (top, dark blue) and Catch*ReSeq (bottom, blue).

D. Clone mean percent spliced in (psi) of *PTPRC* cassette exons A (black) and C (orange) by Catch*ReSeq plotted against clone size (cells per clone, log scale). Each point is one lineage-barcoded clone with ≥3 informative cells, sized by its number of informative cells. Lines are linear fits with 95% confidence bands; rho, Spearman correlation coefficient.

E. Between-cell dispersion of the soluble *IL7R* isoform by method (10X, dark blue; Catch*ReSeq, blue). Per-cell soluble fraction among soluble-*IL7R*+ cells, binned at 0.2 and plotted as a share of each method's soluble+ cells (Catch*ReSeq n = 46, 10X n = 22). Exon inclusion in 10X was quantified using BRIE2.

F. Detection frequency by Catch*ReSeq (blue) or 10X (dark blue) of five ESRP2-spliced exons within ESRP2 knockout cells expressing the parent gene. Each method scored on exon inclusion only for exon-targeting probes (*Kras*, *Lsm14b*, *Dgkd*), inclusion and exclusion for splice-targeting probes (*Nf2*, *Slk*). 10X isoform counts were quantified using BRIE2. Significance calculated using paired McNemar test. Error bars represent 95% Clopper-Pearson confidence intervals. $^*p < 0.05$, $^{**}p < 0.01$, $^{***}p < 0.001$, n.s. = not significant.

G. Cells expressing the five Esrp2 target exons and their genes. Cells per target exon and genotype (WT, KO): 10X total, all cells with ≥1 10X UMI for the gene (gray); 10X, the subset BRIE2 can assign to inclusion or exclusion (dark blue); Ca*Re, cells with ≥1 Catch*ReSeq UMI at the cassette exon (blue). All cells with ≥1 10X gene UMI, pooled across clusters, matching the denominator of Fig. 4M.

H-I. Violin plots comparing Catch*ReSeq counts across cluster (top, cluster 5 highlighted) and by Catch*ReSeq positivity (bottom) in knockout *Kras*+ cells in hepatocyte clusters to total UMI per cell (M) and percent mitochondrial reads (N). Significance given by BH-adjusted Wilcoxon p.

J. *Kras* exon 5 detection rate across hepatocyte clusters in Esrp2 knockout, restricted to *Kras*-expressing cells. Catch*ReSeq+ out of total counts listed above each bar. Dashed line is at the overall rate of 0.013. Omnibus Fisher's exact p = 0.018.

**Table 1: Rhesus macaque specimen information**

| Sample | InTextName | Figs | Monkey | Tissue | DPI | DP. ART | PlasmaSIV (copies/mL) | SortGate |
|---|---|---|---|---|---|---|---|---|
| W0 | uninfected | 1 | HD88 | PBMC | 0 | 0 | N/A | CD3+CD8- |
| D0 | uninfected | 1 | LP29 | PBMC | 0 | 0 | N/A | CD3+CD8- |
| W2 | *in vivo* | 1,3 | A8R095 | PBMC | 14 | 0 | 95,000,000 | CD3+CD8- |
| Pacute | *in vivo* | 1,3 | 08M171 | PBMC | 14 | 0 | 11,000,000 | CD3+CD8- |
| D195 | ART | 1,3 | LP29 | PBMC | 195 | 154 | <15 | CD3+CD8- |
| D13 | *in vivo* | 2 | JK85 | PBMC | 13 | 0 | 93,000,000 | CD3+CD8- |
| Sample | InTextName | Figs | Monkey | Tissue | DPI | DP. ART | %p24+ | SortGate |
| Invitro | *in vitro* | 1,3 | KP81 | PBMC | 7 | 0 | data not available | CD3+CD8- |
| GFP_ CD4 | *in vitro* | 4 | 20-228 | PBMC | 12 | 0 | 1.49 | CD3+CD8- |

**Table 2: PWH and PWoH donor information**

| Sample | PID | HIV status |
|---|---|---|
| S17 | 30832 | HIV- |
| S18 | 41627 | HIV- |
| S19 | 30634 | HIV+ |
| S20 | 44084 | HIV+ |

**Table 3: Catch*ReSeq primers**

All primers are in forward orientation (FWD) unless marked REV

| Name | Sequence |
|---|---|
| 10X homology (REV) | CTACACGACGCTCTTCCGATCT |
| SIV and HIV | |
| SIV D4 | GGCTAATACATCTTCTGCATCAAAC |
| SIV env | CCAGCAGACCCATATCCAACAGG |
| SIV SSenv | GCACCTCCAGGTTATGCTTTGC |
| SIV D1 | AGGAAGAGGCCTCCGGTTGCA |
| SIV pol | TGGAAGCATTTCTCATGGCATTGACA |
| HIV D1 | GCAGGACTCGGCTTGCTG |
| HIV D4 | CGAAACTGACAGAGGATAGATGGAAC |
| HIV pol | CTTTGGAAAGGACCAGCAAAGCT |
| HIV env | GCAGCAGGAAGCACTATGGG |
| HIV nef | TTAAAAGAAAAGGGGGGACTGGA |
| Mouse | |
| Kras | TGGTGAGAGAGATCCGACAGT |
| Nf2 | GCCTTGAAACTCAAAGAGCGGG |
| Slk | CTAACCGCCTGAGAGACGAAG |
| Lsm14b | GAACAGAAGACCTCCACGGAG |
| Dgkd | ATAAGGACATCTTCACGCGGC |
| Macaque | |
| KLRB1 | ACCCAAGGACTCAGGCCCAG |
| PTPRC | TGCTCTTCCGATCTGGCATTTGTCTTTGCC TTTCTGGAC |
| BCL2L1 | TGCTCTTCCGATCTACCTGAATGACCACCT AGAGCC |
| IL7R | TGCTCTTCCGATCTTCTGGAGTGAATGGA GTCCAAGTTA |

**Supplementary Note 1: Frouard et al. HIV-seq did not increase HIV+ cell yield**
An alternative approach to host-virus profiling developed by Frouard et al. adds viral-specific spike-in primers at 10X 5' reverse transcription to improve viral sequence capture[1]. In our reanalysis of the deposited datasets (GSE266455), the addition of capture primers did not increase HIV+ cell frequency, neither in ART-suppressed donors (7 / 17,043 (0.041%) with primers versus 7 / 13,798 (0.051%) without; Fisher's exact p = 0.79) nor in viremic donors (PID1052: 4 / 6,752 versus 2 / 3,377, p = 1.00; PID8027: 90 / 8,908 versus 58 / 7,206, p = 0.18).

**Supplementary Note 2: Internal polyA usage exceeded random under every definition**
To address reads that may be assigned to an internal polyA by chance due to their density in the SIV genome, every parameter set was additionally applied to 20,000 positions drawn uniformly along the 9,602 bp SIVmac239 transcript and classified into internal/terminal/unexplained. Relative to this control, real reads were 1.9x (Read1) and 1.2x (Read2) more likely than chance to end near an internal polyA, and 52x and 10x less likely to end near the terminal polyA site (medians across 33 parameter sets per anchor). Combining both terms, the internal:terminal odds observed in the data exceeded the odds at random positions by a median of 97x (Read1) and 16x (Read2) across every parameter set tested.

**Supplementary Note 3: Replication incompetence did not explain non-canonical splicing**
Proviral genomes may harbor mutations that render them unable to produce replication competent virus although they may be transcribed and even spliced[2]. To determine if the non-canonical splicing we observed was due to provirus mutation and possible replication incompetence, we looked for cells that shared the same non-canonical splicing pattern in all reads aligning to those regions after filtering. Of all cells with non-canonical splicing, no cells had evidence of proviral mutation, including ART-suppressed cells. We cannot rule out proviral defects outside the regions observed by sequencing.

**Supplementary Note 4: Host library size did not explain host program usage**
Infected cells carried more host RNA than uninfected cells in every sample (**Supp Fig 3B, top**), so any program difference could in principle reflect library size rather than cell state. We controlled for size by matching rather than by regression to avoid removing any true biological signal. Uninfected reference cells were drawn to match each infected population's host size distribution over ten quantile bins, up to five reference cells per infected cell, and the same Wilcoxon test was repeated at equal depth without modelling depth (**Methods**). Library size matching left every conclusion in the main text intact (**Supp Fig 3B, bottom**).

**Supplementary Note 5: 10X detection was biased toward high viral transcript load**
Among infected cells recovered by Catch*ReSeq, the 10X library called 100% of cells above 64 viral UMI (99–100% above 32) but only 5.7–15.5% of low viral transcript load cells (2–7 UMI) (**Supp Fig 3C**). Restricting analysis to 10X-detected cells returned essentially the high viral transcript load population alone (69–79% of 10X-detected), such that infected cells appeared depleted of the resting/memory program in both viremic animals (padj < 0.0001) and showed no enrichment *in vitro* (padj 0.82). This contrasts with Catch*ReSeq where *in vitro* infected cells

were enriched for the resting/memory program (log2FC +0.45, padj < 0.0001), a trend carried by the low viral transcript load cells (+0.64, padj < 0.0001).

**Supplementary Note 6: Reanalysis of PWH data replicated published frequencies**
Collora et al.[3] and Frouard et al. datasets were realigned to a joint GRCh38 and HIV-1 HXB2 reference genome. This approach differed from the two-pass approach used by Collora et al. (host-only alignment followed by STAR alignment against autologous proviral sequences with HXB2), and from that of Frouard et al. (joint GRCh38 and subtype B consensus sequence generated from a Los Alamos Group M alignment). Transcriptome QC matched what was used in each study: ≥500 detected genes and ≤10% mitochondrial reads for Collora et al. and ≥500 genes and ≤15% mitochondrial reads for Frouard et al. HIV+ calling threshold matched what was reported for Collora et al. (≥2 HIV-1 UMIs or ≥4 reads of a single UMI), although we raised the threshold of Frouard et al. from their ≥1 HIV-1 read to a minimum of ≥2 HIV-1 reads per cell.

Comparing HIV+ cell recovery from our reanalysis versus the original authors', for Collora et al. we recovered 102 versus their 81 HIV RNA+ cells out of 56,544 versus 52,473 QC-passing CD4+ T cells during viremia (0.180% vs 0.154%), and 10 versus their 9 out of 39,099 versus 36,806 under ART (0.0256% vs 0.0245%). For Frouard et al. we recovered 600 versus their 1,008 out of 63,491 versus 73,382 during viremia (0.945% vs 1.374%), and 9 versus their 25 out of 60,584 versus 75,158 under ART (0.0149% vs 0.0333%). The difference between HIV+ cells in viremia from our reanalysis and Frouard et al. was due to our more stringent HIV+ calling threshold (156 of 408), our more stringent restriction of HIV+ calling to CD4+ T cells (164 of 408), and our alignment to HXB2 rather than their subtype B consensus (88 of 408).

**Supplementary Note 7: Splice+ state versus high viral transcript load is a consequence of detection**
In reanalysis of human datasets, 67.7% of HIV RNA+ cells in Collora et al. and 67.3% in Frouard et al. carried exactly one viral UMI (72.9% and 77.8% carry at most two). Splice status and viral transcript load were nearly interchangeable: viral transcript load classified splice status with ROC AUC 0.990 (95% CI 0.970–1.000) in Collora et al. and 0.978 (0.948–0.994) in Frouard et al., with McFadden pseudo-$R^2$ of 0.77 and 0.70 for logistic splice status on $\log_2$ viral transcript load. Stratifying cells by their exact viral UMI count left essentially no residual variation to explain: in Collora et al. not a single viral transcript load value contains both a splice+ and a splice− cell, and in Frouard et al. only 3 viral transcript load values do (107 cells), retaining 4% of the discordance present between unmatched cells.

**Supplementary Note 8: Barcode and SIV infection were independent**
We confirmed that lentivirally-delivered barcode and SIV infection were independent: SIV-transcribing rates were similar in barcoded vs non-barcoded cells (1.14% vs 1.50%), as were barcoding rates in SIV transcribing vs non-transcribing cells (25.9% vs 31.5%).
